# Metastable Dimeric Intermediate Drives Native-like Spidroin Fibrillogenesis, while Dehydration Promotes Cross-β Transition

**DOI:** 10.64898/2026.09.09.749980

**Authors:** Haruya Kajimoto, Kento Yonezawa, Kok Sim Chan, Kiichi Hayashi, Yusuke Okamoto, Rakuri Aiba, Yuki Nakatani, Kenta Kimura, Minami Bessho, Yoichi Yamazaki, Sachiko Toma-Fukai, Takayuki Uchihashi, Takehiro K. Sato, Hironari Kamikubo

**Affiliations:** Division of Materials Science, Graduate School of Science and Technology, Nara Institute of Science and Technology, 8916-5, Takayama, Ikoma, Nara 630-0192, Japan; Center for Digital Green-innovation, Nara Institute of Science and Technology, 8916-5, Takayama, Ikoma, Nara 630-0192, Japan; Department of Physics, Nagoya University, Furo-cho, Chikusa-ku, Nagoya, Aichi 464-8602, Japan; Spiber inc., 234-1, Kakuganji-mizukami, Tsuruoka, Yamagata, Japan

**Keywords:** Spider silk, Spidroin, Self-assembly, Amyloid, Structural polymorphism

## Abstract

The exceptional mechanical properties of spider dragline silk originate from hierarchical spidroin self-assembly, yet mimicking this process to generate native-like structures remains challenging. Here, we report a metastable dimeric intermediate as an essential molecular factor for fibrillogenesis. Small-angle X-ray scattering (SAXS) and atomic force microscopy (AFM) revealed a compact core stabilized by solvent-excluded “dry interfaces” surrounded by extended random coil regions. Within the core, a subset of the polyalanine blocks is held by side-chain packing rather than by main-chain β-sheets; the majority remain exposed in the surrounding coil regions. Upon lowering the denaturant concentration, these exposed polyalanine segments pair with their counterparts at a fibril end, forming intermolecular β-sheets that act as linkers and drive the stepwise end-to-end polymerization of the dimeric particles. In solution, the resulting fibrils exhibit an ordered β-structure in which the polypeptide chains run parallel to the fibril axis, in contrast to amyloid-like cross-β, in which the strands lie perpendicular to it; this matches key features of native silk nanofibril architecture. However, X-ray diffraction (XRD) analysis demonstrates that these native-like nanofibrils undergo an effectively irreversible transition toward amyloid-like cross-β structures upon drying. Native-like fibrils are thus achievable by self-assembly, but structural control is lost during the final dehydration step. Accordingly, we propose a general principle for decoupling self-assembly from dehydration: native-like nanofibrils are formed first, then dried under conditions that restrict molecular-chain mobility—for example, mechanical constraint or immobilization on a surface—which we expect to suppress thermodynamic relaxation into cross-β structures.

**Significance Statement:** Artificially reproducing the exceptional toughness of spider dragline silk remains a long-standing challenge. Here, we identify a key molecular factor that governs native-like fibrillogenesis. A metastable dimer serves as a key nucleus for fibrillogenesis, forming native-like nanofibrils in solution with an ordered, fiber-axis-parallel β-structure distinct from amyloid-like cross-β. Yet, these nanofibrils transition, effectively irreversibly, into cross-β structures typical of amyloids upon drying. Thus, structural control during dehydration emerges as a previously overlooked bottleneck. Achieving artificial spider silk requires a two-stage design principle that decouples self-assembly from dehydration. Moreover, this perspective of two-stage optimization addresses a challenge of integrally designing self-assembly and final dehydration/solidification processes, not only for spider silk but for protein-based biomaterials in general.

## Introduction

Spider dragline silk (major ampullate [MA] silk) has garnered notable attention as a next-generation structural biopolymer owing to its combination of steel-like tensile strength and extraordinary toughness (1). These exceptional mechanical properties originate from the hierarchical nanostructure formed by spidroins. Notably, the primary structure of spidroins is composed mainly of repetitive sequences of polyalanine (poly-Ala) and glycine-rich segments (2, 3), which self-assemble to form a nanocomposite in which crystalline regions are dispersed within an amorphous matrix. Particularly, β-sheet crystals formed by poly-Ala segments are highly aligned parallel to the fiber axis, and this parallel alignment is considered to be one of the major factors underlying the unique toughness of spider silk (4–10).

Over the past few decades, intensive efforts have been made to create artificial spider silks using recombinant spidroins. However, reproducing structure and properties comparable to those of native silk remains an unresolved challenge. When spidroins self-assemble *in vitro*, thermodynamically stable fibrils containing β-sheets are reported to form (11–17). Where the architecture of these fibrils has been determined by X-ray diffraction, the assignment has been to an amyloid-like cross-β structure, distinct from that of native spider silk (18). It should be noted that such measurements are necessarily made on dried specimens, since the orientation of β-strands relative to the fiber axis cannot be determined by methods applicable in solution; the cross-β architecture has subsequently been assumed to be present in solution as well. Throughout this paper we distinguish the two architectures by the orientation of the polypeptide chain relative to the fiber axis. In a cross-β architecture the β-strands lie perpendicular to the axis, the inter-strand hydrogen bonds run parallel to it, and the axial repeat is the inter-strand spacing of ∼0.47 nm; the 0.47 nm reflection is accordingly meridional. In native dragline silk the chains instead run parallel to the fiber axis, so that the same reflection appears on the equator and the axial repeat is set by the length of the extended molecule. We refer to the latter as the fiber-axis-parallel architecture. We use fiber axis for the macroscopic fiber and fibril axis for the nanoscale fibril; the term fiber-axis-parallel denotes the architecture itself and applies at either scale. This distinction concerns orientation alone and is independent of hydrogen-bond register, which we specify separately as parallel or antiparallel. The formation of a cross-β architecture rather than the fiber-axis-parallel architecture of native silk is considered one of the major reasons why artificial silks fail to exhibit sufficient toughness and strength. Yet, the molecular factors that render the pathway to the native structure inaccessible *in vitro* remain unelucidated (10).

In the *in vivo* spinning duct, environmental factors such as pH gradients, changes in ion composition, and dehydration precisely control structural transitions of spidroins (19–23). Indeed, fibrils have been suggested to exist in the middle section of the major ampullate (MA) gland sac (24, 25), and dragline silk itself is reported to possess a hierarchical structure composed of bundled fibrils (26–30). While precursor processes such as micelle formation, liquid–liquid phase separation (LLPS), and the formation of liquid-crystalline phases have been proposed (23, 31–35), the molecular mechanisms underlying fibrillogenesis—specifically, what structural unit initiates nucleation and how it elongates—remain elusive. Consequently, the correspondence between these *in vivo* fibrils and those obtained *in vitro* remains unclear, and the essential process required to avoid amyloid-like aggregation (cross-β) and guide the system toward a native-like orientation in hydrated states has not been identified.

Dimerization of spidroins is itself well documented, but in a different structural context. The N-terminal domain (NTD) forms a pH- and ion-dependent dimer whose mechanism is conserved across silk types (36–40), and the C-terminal domain forms a constitutive dimer (41); consistent with this, mass photometry indicates that spidroin in native major ampullate dope is predominantly dimeric (42). These terminal-domain dimers act as regulatory devices: they maintain solubility during storage and trigger assembly upon acidification. Whether the repetitive region itself can form a defined dimer, and whether such a species would participate in fibril growth rather than merely regulate it, has not been examined. This question is central to the present work, because the construct used here consists solely of repetitive sequence and carries no terminal domain.

In this study, to elucidate the molecular-level factors governing the pathway to the native structure, we explored kinetic intermediates in the spidroin folding pathway—specifically, during the reconstitution process from a fully denatured state. We identified a “metastable dimer” that functions as a key nucleus for fibrillogenesis. Detailed structural analyses revealed that this dimer is stabilized by the formation of intermolecular solvent-excluded interfaces. We demonstrate that fibril elongation proceeds via stepwise end-to-end polymerization of these dimers as building units, resulting in the uniaxial growth of fibrils that, in solution, exhibit an ordered, fiber-axis-parallel β-structure distinct from amyloid-like cross-β and consistent with key features of native silk nanofibril architecture. Furthermore, we observed an effectively irreversible transition of these “native-like fibrils” into a cross-β structure during the drying process. These findings clarify that the lack of structural control during the final dehydration step, rather than the self-assembly process itself, is the primary bottleneck underlying the failure of conventional reconstitution strategies.

## Results

### Identification of a Fibrillogenic Intermediate

First, we evaluated the solubility behavior of spidroin as a function of denaturant concentration. Purified spidroin powder exhibited high solubility (>20 mg/mL) in aqueous urea solutions of 6 M or higher, whereas aggregation increased sharply at 4 M or lower concentrations (Fig. S1). Next, to identify the structural state competent for fibrillogenesis, we prepared two samples with different solvation histories: “UU,” prepared by dissolving purified powder directly in 6 M urea, and “GU,” prepared by first dissolving the powder in 5 M guanidinium thiocyanate (GdmSCN), a strong denaturant, to fully unfold the structure, followed by buffer exchange into 6 M urea.

We diluted these samples to conditions where aggregation increases (3 M urea) and tracked the reaction using a Thioflavin T (ThT) assay. In 6 M urea, both UU and GU showed low, time-invariant ThT fluorescence (Fig. 1A). Upon dilution to 3 M urea, GU showed no change in fluorescence intensity. In contrast, UU exhibited a marked increase in fluorescence intensity, reaching saturation by approximately 40 h (Fig. 1B).

**Fig. 1.**
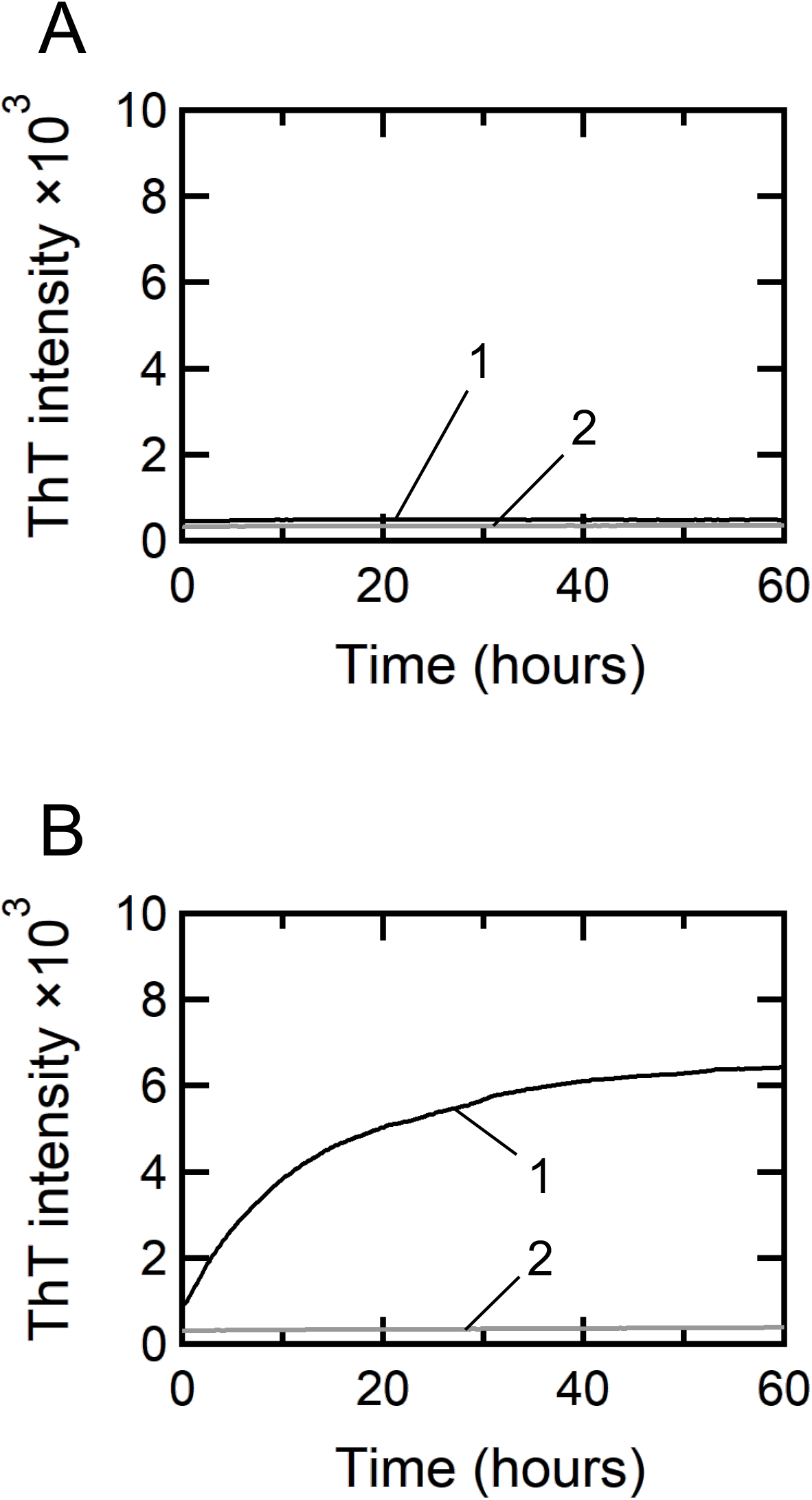
Time course of fibrillogenesis monitored by ThT fluorescence. (*A*) ThT fluorescence in 6 M urea, recorded for 60 h immediately after preparing UU (urea–urea; Line 1) and GU (guanidine–urea; Line 2). Both remained low and time-invariant. (*B*) ThT fluorescence in 3 M urea, recorded from immediately after dilution. UU (Line 1) increased with time and saturated within several tens of hours, whereas GU (Line 2) showed no distinct change, indicating that only UU retains fibrillogenic competence. ThT, thioflavin T.

Consistently, when samples 60 h post-dilution were observed by atomic force microscopy (AFM), no distinct structures were seen in GU, whereas UU contained fibrils with uniform widths and lengths reaching the micrometer scale (Fig. 2 A and B). These fibrils possessed periodic corrugation with a pitch of ∼31.3 nm (Fig. 2C). N-terminal labeling experiments using Ni-NTA-functionalized gold nanoparticles yielded an inter-particle distance of approximately 27.4 nm (Fig. 2D). As these values correspond to the estimated axial length of a spidroin molecule, this suggests that molecular chains adopt an extended, end-to-end registry along the fibril axis (discussed below).

**Fig. 2.**
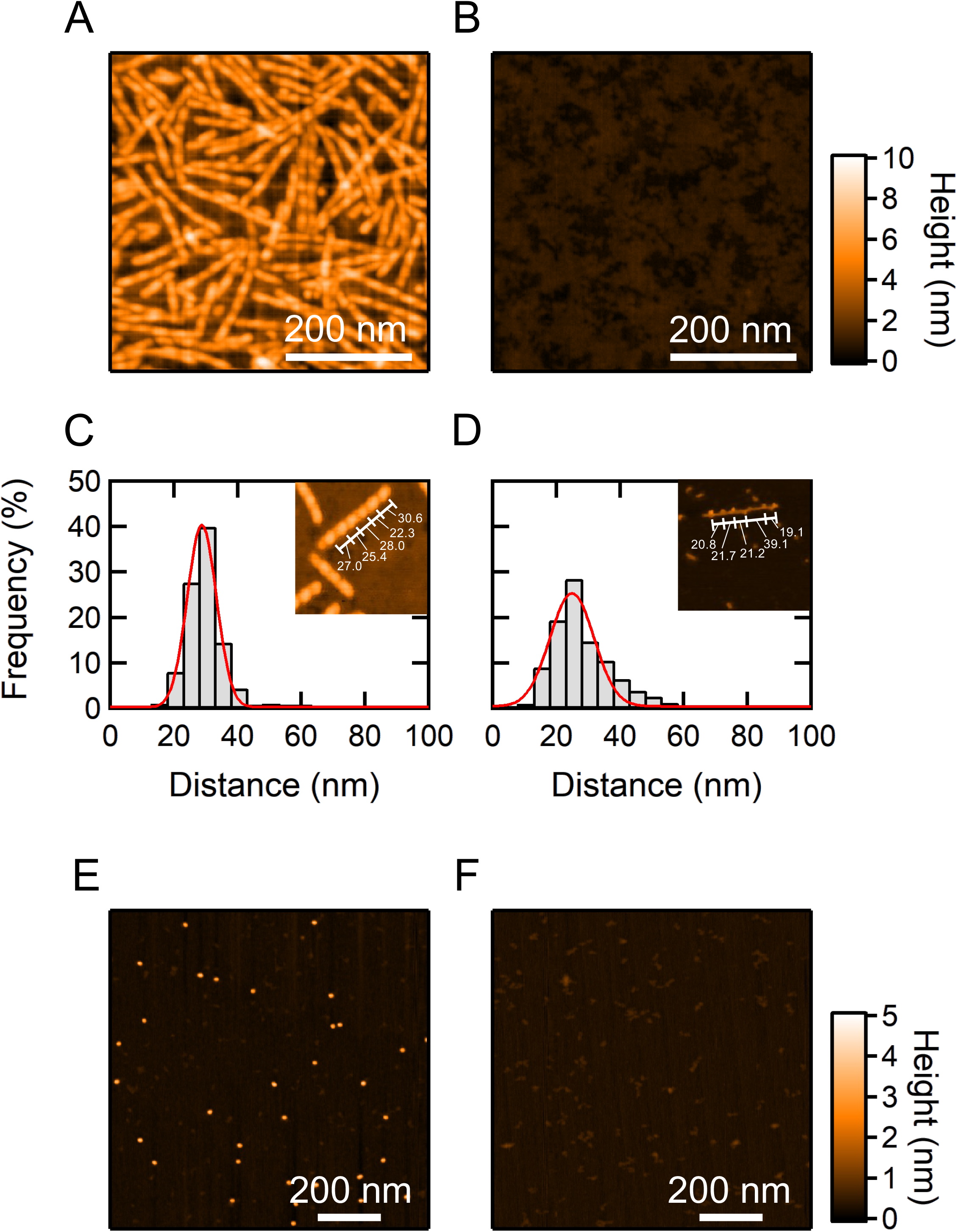
Morphological evaluation of UU and GU by AFM. (*A* and *B*) AFM images after incubation for 60 h under 3 M urea conditions ((*A*) UU; (*B*) GU). No distinct higher-order structures were observed in GU, whereas UU contained numerous fibrils of nearly uniform width, reaching micrometer lengths and showing a periodic height modulation along the fibril axis. (*C*) Distribution of the distance between adjacent protrusions in the periodic height profile along UU fibrils; the red line is a Gaussian fit, giving an average interval of 31.3 nm. (*D*) Distribution of the distance between adjacent gold nanoparticles in fibrils labeled at the N-terminal His-tag with Ni-NTA-functionalized gold nanoparticles; the average interval was 27.4 nm. (*E* and *F*) AFM images of UU (*E*) and GU (*F*) under 6 M urea conditions, prior to dilution to 3 M urea. No significant particulate structures were seen in GU, whereas UU contained numerous dispersed nanoparticles of height approximately 2.6 nm. Scale bars, 200 nm. UU, urea–urea; GU, guanidine–urea; AFM, atomic force microscopy.

To identify structural features governing fibrillogenic competence, we observed the initial state (in 6 M urea) by AFM. While GU did not show distinct images, UU contained numerous dispersed “nanoparticles” with a height of approximately 2.6 nm (Fig. 2 E and F). This difference indicates that the nanoparticles were likely formed during the powder preparation process—specifically via ethanol precipitation from GdmSCN solution (see Materials and Methods). These pre-formed nanoparticles thus persisted in UU but were completely dissociated upon re-exposure to GdmSCN in GU. The results demonstrate that this nanoparticle is the physical entity responsible for fibrillogenic competence after dilution and serves as a key precursor for growth into fibrils.

### Structural Characterization of the Metastable Dimer

To quantify the structural characteristics of the precursor nanoparticles, we performed small-angle X-ray scattering (SAXS) measurements on UU and GU in 6 M urea solution. The forward scattering intensity normalized by concentration, *I*(0)/*Conc.*, calculated from Guinier analysis (Fig. 3 A and B; Fig. S2), was approximately twice as high for UU (1.12 × 10^−3^) as for GU (5.88 × 10^−4^) (Table 1). Because *I*(0)/*Conc.* is proportional to the molecular weight of the scatterer, this reveals that while GU exists as a monomer, UU forms a dimer. The same comparison made using the forward scattering intensity obtained from the distance distribution function P(r), which draws on the scattering curve over a wide angular range rather than on the low-angle region alone, gives the same result (Fig. S2 C and D).

**Fig. 3.**
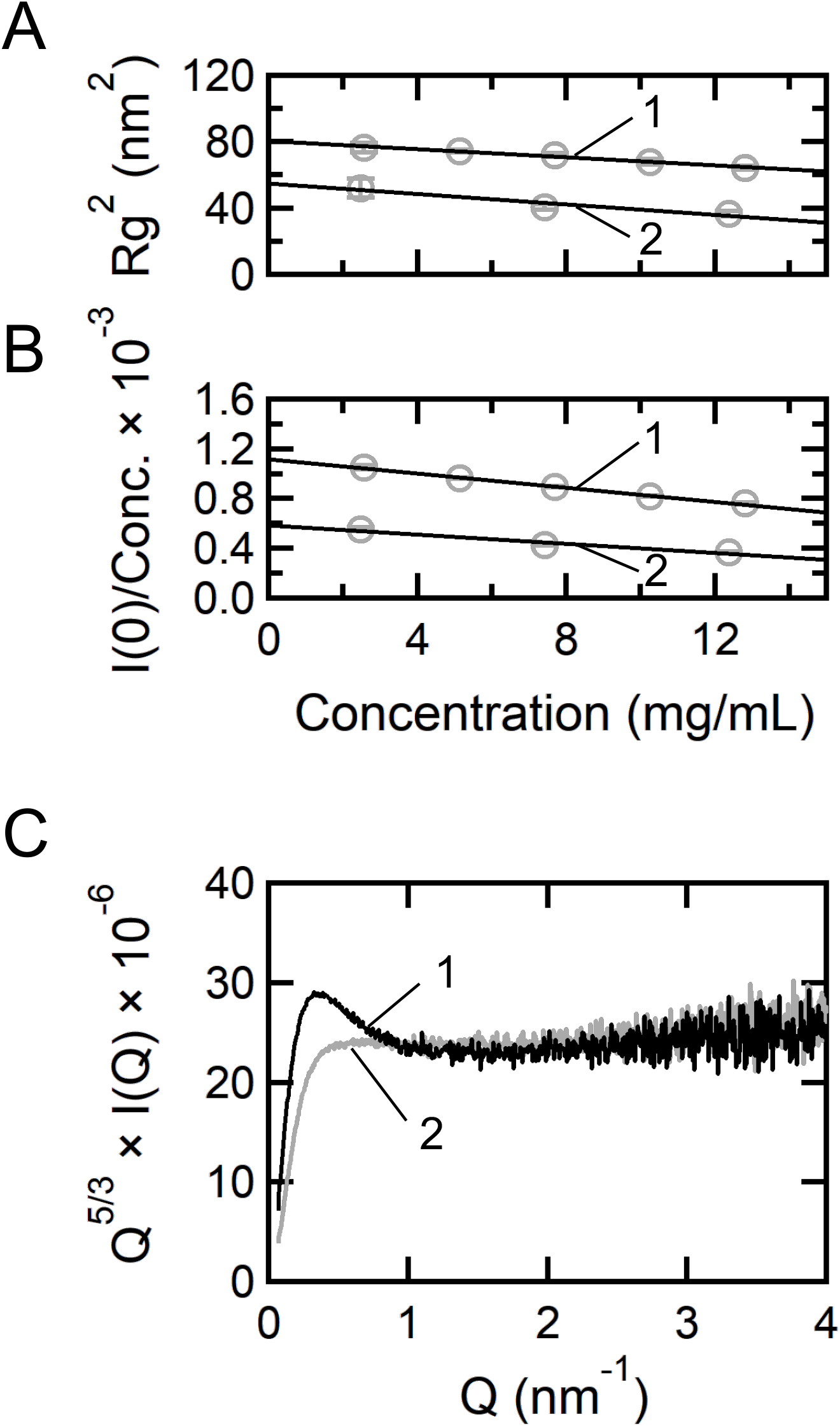
Small-angle X-ray scattering of UU and GU in 6 M urea. (*A* and *B*) Concentration dependence of the radius of gyration *R_g_* (*A*) and of the concentration-normalized forward scattering intensity *I*(0)/*Conc.* (*B*), from Guinier analysis. For both UU (Line 1) and GU (Line 2) the values decreased slightly with concentration because of inter-particle interference; values extrapolated to zero concentration were used (Table 1). (*C*) Kratky plots. GU (Line 2) showed the plateau typical of a random coil, whereas UU (Line 1) showed a broad peak near *Q* ∼ 0.32 nm⁻¹, indicating a partially compact core. SAXS, small-angle X-ray scattering; *R_g_*, radius of gyration; *I*(0), forward scattering intensity; *Q*, scattering vector; UU, urea–urea; GU, guanidine–urea.

**Table 1.** Concentration-normalized forward scattering intensity and radius of gyration values for urea–urea and guanidine–urea obtained from Guinier analysis.

|  | *I*(0)/*Conc*.<sup>a</sup> |  | *R<sub>g</sub>* (nm)<sup>a</sup> |  |
|---|---|---|---|---|
| UU | 1.12×10<sup>−3</sup> | ±1.02×10<sup>−5</sup> | 8.97 | ±0.95 |
| GU | 5.88×10<sup>−4</sup> | ±3.14×10<sup>−5</sup> | 7.40 | ±1.91 |

<sup>a</sup> Values extrapolated to zero concentration were adopted to eliminate the effects of inter-particle interference.
*I*(0), forward scattering intensity; *Conc.*, concentration; *R<sub>g</sub>*, radius of gyration; UU, urea–urea; GU, guanidine–urea.

To assess particle shape, we constructed Kratky plots (Fig. 3C; Fig. S3). GU exhibited the behavior typical of a random coil, reaching a plateau with increasing *Q*. UU showed the same plateau at wide angles but, in addition, a broad peak in the low-angle region (*Q* ∼ 0.32 nm⁻¹). The scattering data therefore establish three points: the particle mass is twice that of GU; a region of the chain is compact rather than coil-like, giving rise to the low-angle peak; and the remainder of the chain behaves as an excluded-volume coil that is indistinguishable from GU. This combination — a compact region surrounded by expanded coils — corresponds to the scattering signature of a branched architecture in which a compact centre is surrounded by expanded chains, rather than to that of either a uniformly globular particle or a fully expanded chain (Fig. S3). We therefore describe the nanoparticle as a dimer comprising a compact core and a surrounding random-coil corona.

Assigning that compact core to the inter-protomer interface does not follow from the scattering data alone; it is an inference, supported by the thermal, spectroscopic, and preparative observations described below and in the Discussion.

Next, we determined whether this dimer exists in a thermodynamically stable equilibrium state or a kinetically trapped metastable state. When SAXS measurements were performed while increasing the temperature under 6 M urea conditions, a cooperative decrease in both *I*(0)/*Conc.* and the radius of gyration (*R_g_*) was observed between 40 °C and 60 °C, indicating the dissociation of dimers into monomers (Fig. 4; Fig. S4). Importantly, the original dimeric structure was not recovered upon cooling the dissociated sample (irreversible denaturation). This fact strongly supports the notion that the UU dimer is not a product of an equilibrium reaction but is a “kinetically trapped metastable state” likely formed during ethanol precipitation following GdmSCN dissolution in the course of powder preparation.

**Fig. 4.**
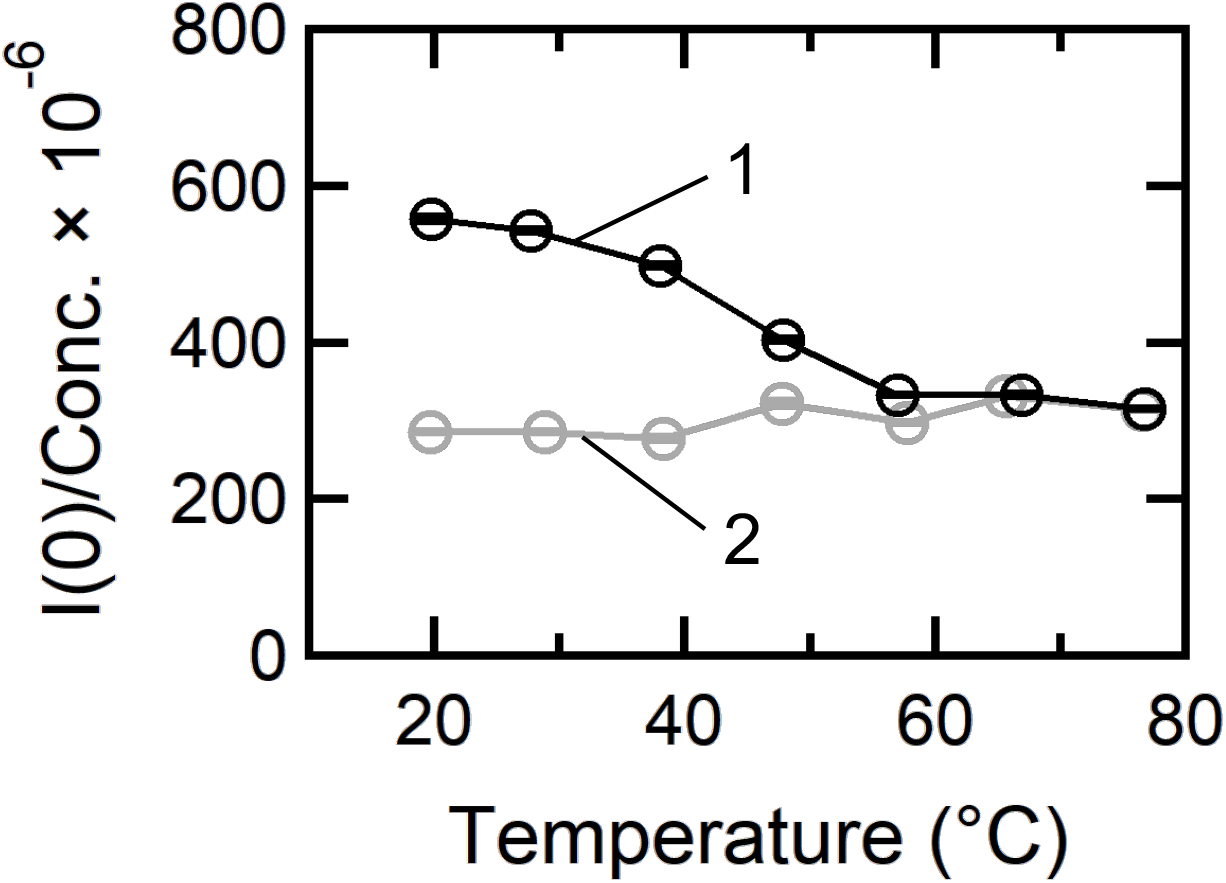
Thermal stability of the UU dimer. *I*(0)/*Conc.* in 6 M urea during heating from 20 °C to 80 °C (Line 1) and cooling from 80 °C to 20 °C (Line 2). The intensity decreased sharply between 40 °C and 60 °C on heating and did not recover on cooling, indicating irreversible dissociation of the UU dimer into monomers. *I*(0), forward scattering intensity; UU, urea–urea.

We use the term metastable here in a specific, operational sense, defined by three observations. First, the dimer is kinetically persistent: in 6 M urea at 27 °C the ThT signal is invariant for at least 60 h (Fig. 1A) and the SAXS parameters do not change over the measurement period, so the species neither dissociates nor polymerizes on this timescale. Second, it is separated from the monomer by an activation barrier, as shown by the irreversibility of thermal dissociation described above. Third, its stability is solvent-dependent: it is stably dispersed in 6 M urea, completely dissociated in 5 M GdmSCN — which is how the monomeric GU reference is prepared — and, at 3 M urea or below, converts into fibrils (Figs. 1B, 5). Metastability therefore denotes here a state that is not the free-energy minimum under the conditions examined and that is separated from the monomer by a kinetic barrier.

**Fig. 5.**
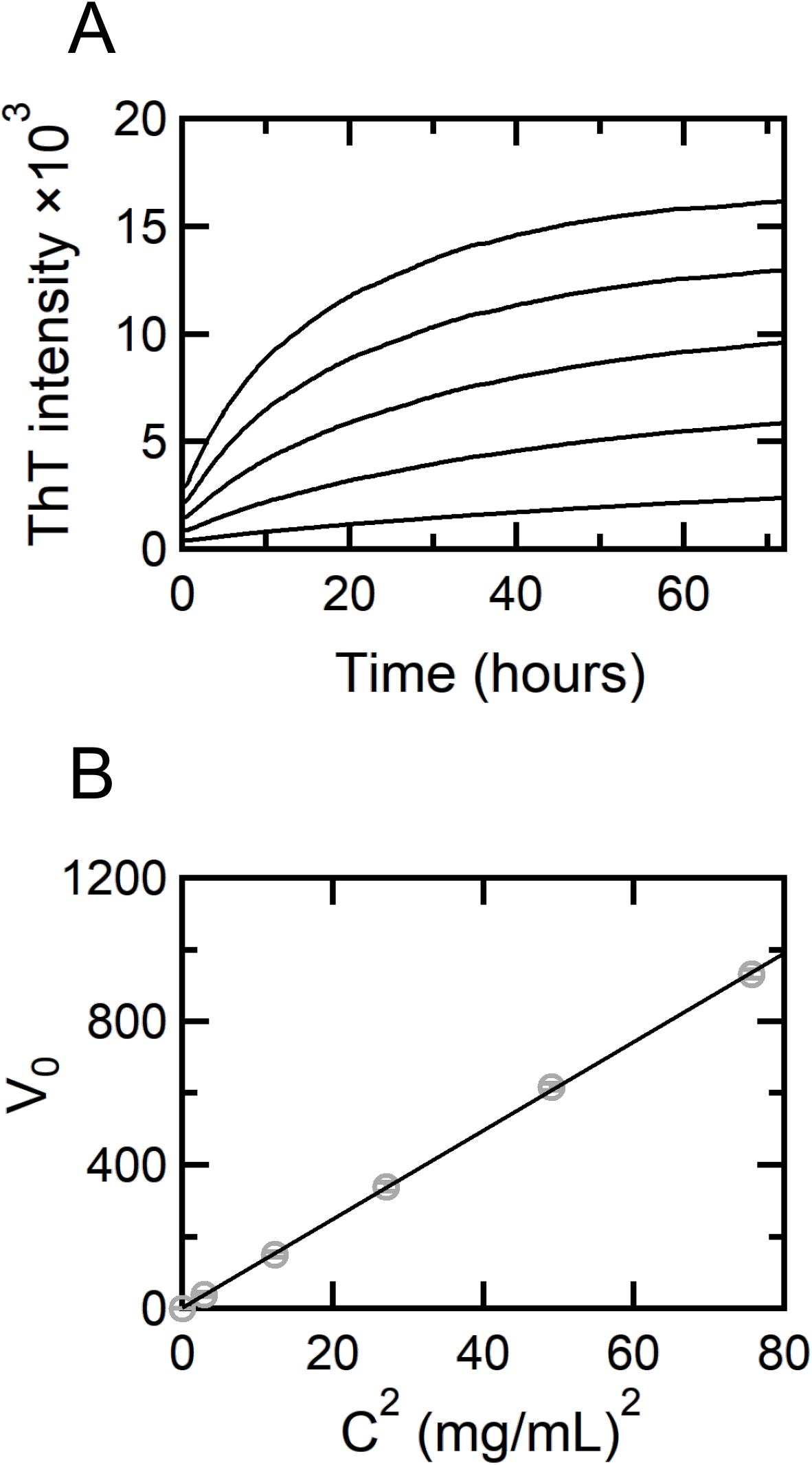
Kinetics of fibril formation for UU under 3 M urea conditions. (*A*) ThT fluorescence at different spidroin concentrations. Higher concentrations gave a faster rise and a higher final intensity. (*B*) Initial velocity of ThT fluorescence (*v*_0_) plotted against the square of the spidroin concentration (*C*^2^). The linear relationship indicates that the initial stage of fibril elongation is rate-limited by a second-order reaction involving collisions between dimers. UU, urea–urea; ThT, thioflavin T; *v*_0_, initial velocity; *C*^2^, square of spidroin concentration.

Circular dichroism (CD) spectroscopy detected no change in secondary structure between UU and GU, both being approximately 70% unstructured (Fig. S5; Table S1). This is not in conflict with the SAXS results, which report on particle size and shape rather than on main-chain hydrogen bonding; the two methods probe different structural levels. Rather, the absence of a CD change is itself informative: it indicates that formation and stabilization of the dimer do not involve the main-chain hydrogen bonding that constitutes β-sheets, and that dimerization instead depends on side-chain interactions. We note that, given the small fraction of residues expected to constitute the core and the restricted spectral window accessible in 6 M urea, a change of this magnitude would not be expected to be resolved by CD.

The construct contains twelve poly-Ala repeats, and the scattering data indicate that only a limited fraction of the chain is compact (Fig. 3C). Only a subset of these repeats can therefore be accommodated within the core; the majority remain within the surrounding random-coil corona, where they are available for intermolecular β-sheet formation once the denaturant concentration is lowered.

### Mechanism of Fibril Elongation

To elucidate the initial process by which dimeric nanoparticles initiate fibril formation, we prepared UU at various concentrations and measured the change in ThT fluorescence intensity immediately after dilution into 3 M urea (Fig. 5A). Higher spidroin concentrations yielded faster initial velocities and increased final fluorescence intensities. A plot of the initial velocity of fluorescence intensity *v*_0_ against the square of the spidroin concentration (*C*^2^) yielded a strong linear relationship (Fig. 5B). Given that the fundamental unit of UU is a dimer, this result suggests that the initial stage of fibril elongation is rate-limited by a second-order reaction involving dimer-dimer collisions.

Next, we investigated whether this dimer functions as a “nucleus” for fibril growth. We prepared UU alone, GU alone, and a mixture of UU and GU (UU:GU = 2:1) and tracked the reaction (Fig. S6). Although the fluorescence intensity of the mixture was higher than the simple sum of UU alone and GU alone, it did not reach the levels expected if GU (monomers) were fully incorporated into elongating fibrils. This indicates that while the UU dimer functions as a nucleus and can promote monomer incorporation, the conversion efficiency is limited. Combined with the kinetic analysis (second-order reaction between dimers), these results indicate that fibril elongation proceeds preferentially via polymerization of dimers rather than the sequential addition of monomers.

To capture structural changes during this elongation process, we performed time-resolved SAXS and CD measurements. In the SAXS profiles (Fig. 6), the peak observed at *Q* ∼ 0.32 nm^−1^ immediately after reaction initiation shifted to the low-angle side (*Q* ∼ 0.20 nm^−1^) in the early stages; subsequently, the intensity increased while the peak position remained unchanged. This indicates uniaxial elongation of cylindrical particles. Concurrently, changes in the shape of CD spectra (Fig. S7) were observed as the reaction proceeded. Secondary structure analysis estimated that unstructured components decreased from 65.0% to 51.4%, and total β-sheet content increased from 19.8% to 28.2% (Fig. S7 B and C; Table S2). Analysis using BeStSel, which distinguishes β-sheet types, identified the formed β-sheets as antiparallel (content: 33%) (Fig. S7 D and E; Table S3).

**Fig. 6.**
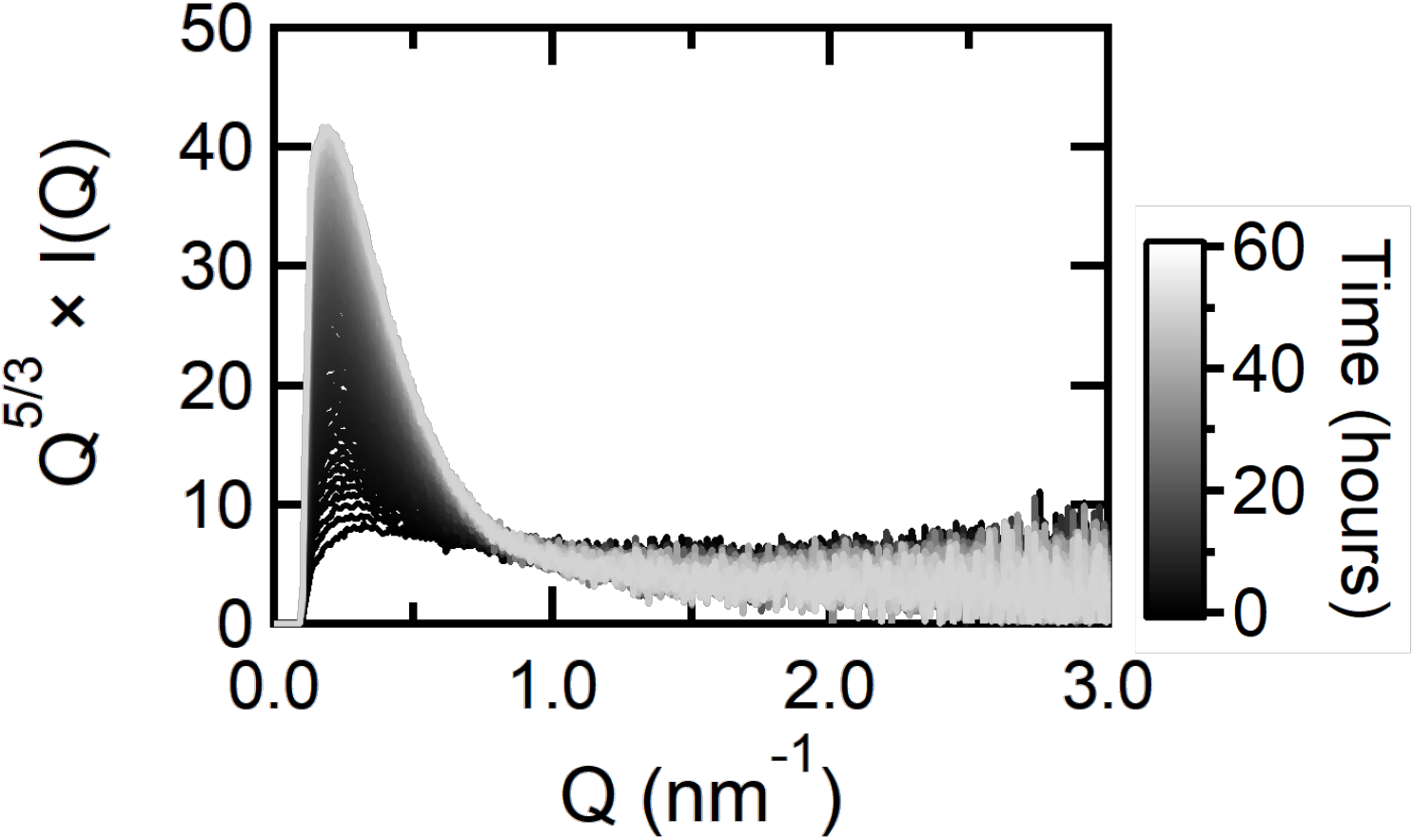
Time-resolved SAXS during fibril elongation. Kratky plots recorded from immediately after diluting UU to 3 M urea up to 60 h. In the initial phase the peak at *Q* ∼ 0.32 nm⁻¹ shifted transiently to *Q* ∼ 0.20 nm⁻¹, after which the intensity increased while the peak position remained unchanged, reflecting uniaxial elongation of cylindrical particles. SAXS, small-angle X-ray scattering; UU, urea–urea; *Q*, scattering vector.

Furthermore, in situ observation by high-speed AFM (Movie S1; Fig. S8) captured the elongation of fibrils at both ends. Detailed image analysis revealed that the fibril length exhibited repeated cycles of an abrupt increase followed by a gradual extension (Fig. S8C). Based on these results, we conclude that fibril elongation is a stepwise addition polymerization of dimeric nanoparticles. Moreover, integrating this with the finding from the preceding section that “dimers possess random coil regions containing poly-Ala segments,” we propose that the elementary process of elongation proceeds as follows: first, the random coil regions of the dimer bind to the random coils at the fibril end to form an “encounter complex,” and subsequently, the β-sheet structure matures over a certain period of time.

### Structural Transition upon Dehydration

Finally, we analyzed the solid-state structure of the obtained fibrils. UU was fibrillized by dialysis against 3 M urea, followed by dialysis against ultrapure water to yield a hydrogel-like sample (Fig. 7A). A solid sample was prepared by drying this gel at room temperature with both ends fixed (Fig. 7B). X-ray diffraction (XRD) measurements of the dried sample revealed distinct anisotropy in the two-dimensional diffraction pattern, indicating that the fibrils were highly oriented in the stretching direction (Fig. 7C). This orientation likely arose from stress applied to the fibrils toward both fixed ends as the hydrogel shrank during drying.

**Fig. 7.**
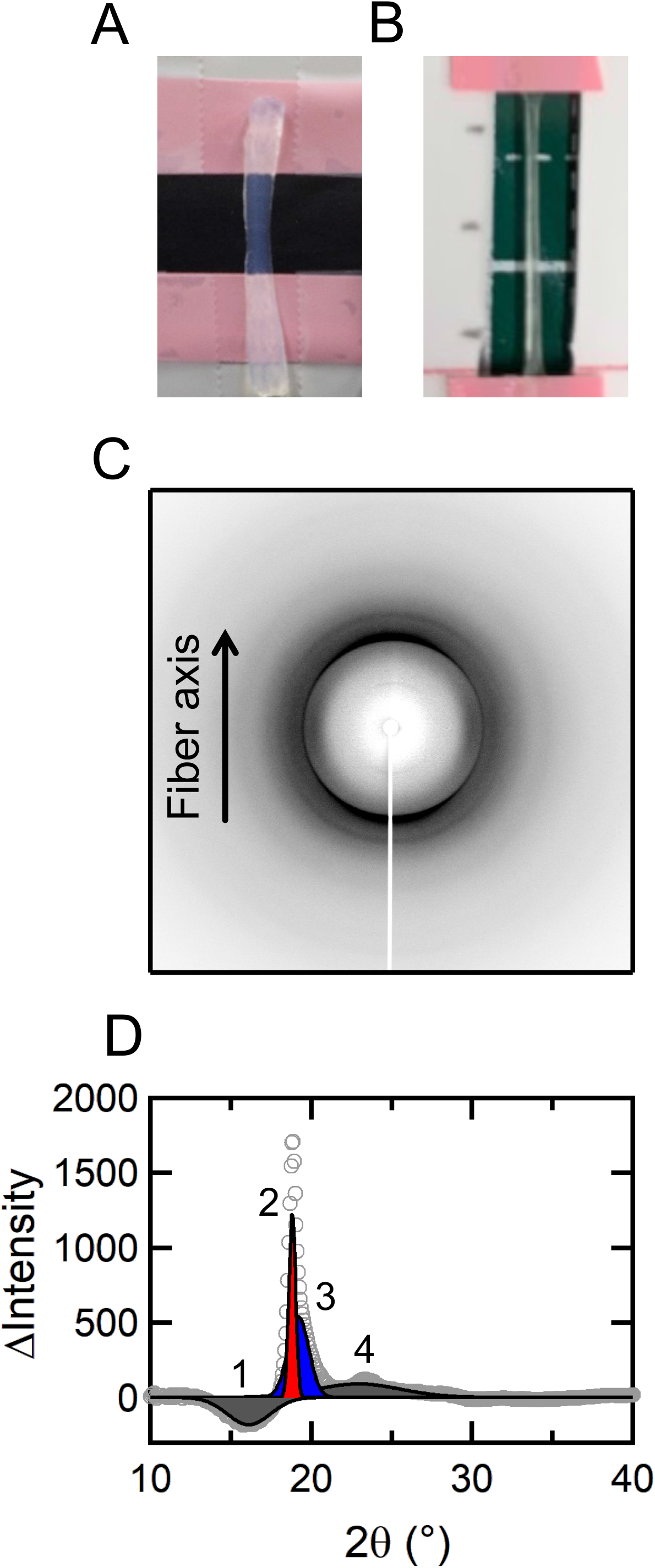
Structural analysis of stretched and dried fibril hydrogels. (*A* and *B*) Photographs of the hydrogel prepared from UU (A) and of the solid sample dried with both ends fixed (*B*). (*C*) Two-dimensional X-ray diffraction image of the dried sample, showing the anisotropy produced by orientation of the fibrils along the fiber axis under the shrinkage stress of drying. (*D*) One-dimensional difference profile (meridional minus equatorial, which removes the isotropic amorphous scattering) and its peak deconvolution. The sharp peaks near 0.47 nm (Peaks 2 and 3) correspond to the inter-strand hydrogen-bonding distance and the broad peak near 0.55 nm (Peak 1) to the inter-sheet stacking distance, as expected for a cross-β structure. UU, urea–urea.

However, the diffraction pattern differed markedly from that of native dragline silk. Sharp reflections corresponding to the inter-strand hydrogen bonding distance (*d* ∼ 0.47 nm) were observed along the meridional direction, and broad reflections corresponding to the inter-sheet stacking distance (*d* ∼ 0.55 nm) appeared along the equatorial direction (Fig. 7D; Table S4; Fig. S9). This pattern is characteristic of a typical “cross-β structure,” where β-strands are arranged perpendicular to the fiber axis.

The aforementioned AFM observations (Fig. 2) and SAXS/CD results (Fig. 6; Fig. S7) suggested that the fibrils in solution possess an ordered, fiber-axis-parallel β-structure distinct from amyloid-like cross-β, consistent with key features of native silk nanofibril architecture (discussed below). Nevertheless, the XRD data after drying indicate a complete cross-β structure. Together, these results lead to the conclusion that native-like nanofibrils formed in solution underwent a dramatic internal structural reorganization during the drying (dehydration) process, transitioning, effectively irreversibly, into a thermodynamically more stable amyloid-like cross-β structure. This provides strong evidence that a major barrier to reproducing the physical properties of spider silk *in vitro* lies not in fibril formation itself, but in the subsequent dehydration and solidification process.

## Discussion

### Formation Mechanism of the Metastable Dimer

The metastable dimer identified in this study functions as a major element initiating fibrillogenesis. The fact that this dimer is not a product of thermodynamic equilibrium but a “kinetically trapped” state formed during the ethanol precipitation process is evident from its irreversible dissociation into monomers upon heating (Fig. 4). Why is this specific association state formed by alcohol treatment? We believe the key lies in two factors: the water activity (*a_w_*) of the solvent and the presence of the strong denaturant GdmSCN.

Typically, irreversible aggregation of proteins occurs via the exclusion of water molecules from contact interfaces. Herein, when ethanol (EtOH) was added stepwise to spidroin in 5 M GdmSCN solution, precipitation was first observed at 75 vol% EtOH. At this point, the initial aqueous solution was diluted four-fold, reducing the GdmSCN concentration to approximately 1.25 M. It is known that *a_w_* drops sharply when EtOH concentration exceeds 60 vol% and decreases to approximately 0.74 at 75 vol% EtOH (43); at this concentration, it is presumed that hydration water on the spidroin surface, particularly in the poly-Ala regions, was largely stripped away. Under such low-*a_w_* conditions, spidroin molecules can overcome the hydration barrier and associate by creating “dry interfaces” where side chains interdigitate tightly—such as packing similar to the Ala-zipper (Fig. 8A) (44–46). Consistent with this, Asakura et al. identified two zipper structures, rectangular and staggered, by XRD and solid-state NMR using poly-Ala peptides of different lengths (45), and demonstrated the coexistence of these arrangements in the poly-Ala region of native dragline silk (46). These facts indicate that poly-Ala regions possess the property of binding by forming dry interfaces.

**Fig. 8.**
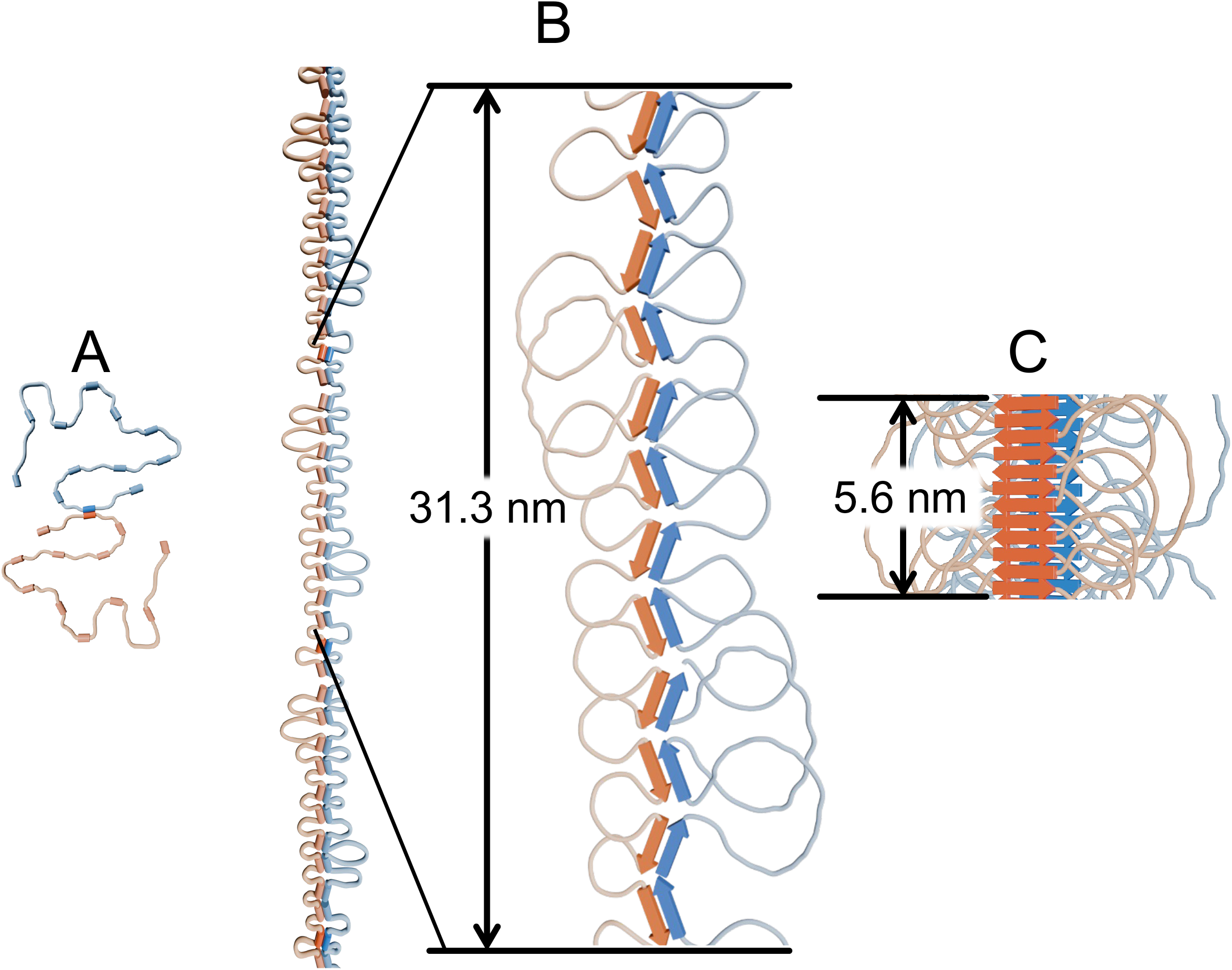
Molecular model of fibrillogenesis mediated by metastable dimers and the dehydration-induced structural transition. (*A*) The metastable dimer: a compact core stabilized by solvent-excluded “dry interfaces,” surrounded by extended random coil regions. A subset of the poly-Ala repeats is held within the core by side-chain packing and does not form main-chain β-sheets; the majority remain exposed in the corona and retain the capacity to form intermolecular β-sheets once the denaturant concentration is lowered. (*B*) Native-like fibril elongation in solution. As the urea concentration falls, the exposed poly-Ala segments form intermolecular β-sheets that act as linkers, driving stepwise addition polymerization of dimer units and uniaxial growth of nanofibrils with a 31.3 nm periodicity—the length of a single extended spidroin molecule—and antiparallel β-sheets running parallel to the fibril axis. (*C*) Transition to cross-β upon dehydration. Color indicates the bonding mode: in solution (*B*) hydrogen bonds form between different strands (alternating blue and orange), whereas in the dried state (*C*) identical strands stack together (same colors). The 5.6 nm dimension is the length expected for a dense cross-β stack of the 12 repeats (0.47 nm × 12).

Crucially, this dehydration occurs in the presence of a high concentration of GdmSCN (≳1 M). GdmSCN strongly inhibits main-chain hydrogen bonding, suppressing the development of higher-order structures including native folds and β-sheets. Thus, while hydration water is largely removed by 75% EtOH, the extensive β-sheet network between main chains remains strongly disfavored by GdmSCN. Consequently, disordered stacking of β-sheets and higher-order oligomerization, which are problematic in typical “dry interface” associations, are suppressed, and only specific side-chain packing is likely to be selectively stabilized. As a result, spidroin molecules under these conditions are placed in a unique energy landscape where “hydration water is largely removed, but long-range β-sheet formation is inhibited,” suggesting that association is arrested at the dimer level stabilized by side-chain packing.

This perspective of association control by a combination of water activity and strong denaturant also accounts for discrepancies with previous work. Scheibel et al. induced fibrillogenesis with high concentrations of potassium phosphate but observed no clear fibril formation for ADF3, the sequence used here, whereas ADF4, which has more poly-Ala repeats and a higher aggregation propensity, did form fibrils after a distinct lag phase (16, 17). Under phosphate conditions the water activity remains high (*a_w_*∼ 0.98) (47), so the dehydration driving force is insufficient and nucleation occurs only as a chance event. With ethanol precipitation (*a_w_* ∼ 0.74), hydration water around the poly-Ala regions is excluded even for ADF3, and nuclei form immediately rather than by chance.

Adjusting ionic strength or salting-out conditions alone is therefore insufficient. What matters is an active dehydration design that controls two parameters at once: how far the water activity is lowered, and how much main-chain folding is permitted while it is.

The metastable dimer thus formed is stably dispersed in 6 M urea but spontaneously initiates polymerization and grows into fibrils when the urea concentration is lowered. This “reaction switch” appears to be precisely governed by the solvent environment. As indicated by SAXS and CD analyses, within the dimeric nanoparticle, dimerization occurs at the core region, while the remaining regions adopt a random-coil (unstructured) conformation (Fig. 8A). These disordered regions inherently contain at least some poly-Ala segments with high β-sheet-forming propensity. High concentrations (6 M) of urea strongly inhibit hydrogen bond formation between protein main chains, keeping these poly-Ala-containing regions in a disordered state with few intermolecular interactions.

However, upon dilution, the urea concentration decreases, weakening the shielding effect on the main chain and enabling the formation of intermolecular hydrogen bonds between exposed random-coil regions. As a result, we propose that dimers are preferentially linked via these random-coil parts acting as “linkers,” undergoing uniaxial stepwise polymerization accompanied by β-sheet formation (Fig. 8B). In essence, urea functions as a “reaction inhibitor” that suppresses main-chain aggregation without destroying the dimer core (side-chain packing), and the release of this inhibition itself serves as the direct trigger driving fibril elongation.

### Structural Characteristics of Fibrils in Solution

The fibrils obtained in this study exhibited drastically different structural characteristics in solution versus in the dry state. Understanding this “structural polymorphism” is key to controlling the physical properties of artificial spider silk.

First, we examined the fibril structure in solution. The periodic corrugation with a pitch of approximately 31.3 nm observed by AFM (Fig. 2C) corresponds well to the scale of a single spidroin molecule. The spidroin used in this study has 12 repeating units, with one unit comprising approximately 35 residues. If the molecule were fully extended, the total length would reach approximately 147 nm, which is much longer than the periodic length observed by AFM. However, in reality, the glycine-rich regions are thought to be contracted as random coils.

This estimate is also consistent with the secondary structure content calculated from CD measurements. The β-sheet content of the fibrils was estimated to be approximately 20–30% (Table S2; Table S3), which agrees well with the assumption that only the regions containing poly-Ala (about 8–10 residues) in each repeat form β-sheets. Calculating the total length when 12 of these β-structure segments (poly-Ala-derived, about 2.8 nm each) are connected in series yields approximately 33.6 nm, which is in excellent agreement with the periodic length observed by AFM (31.3 nm). Furthermore, the distance between gold nanoparticles bound to His-tags at the N-terminus—specifically, the inter-N-terminal distance of spidroins aligned on the fibril (approx. 27.4 nm) (Fig. 2D)—also supports this value.

Collectively, these findings support a model in which the β-structure within spidroin molecules is organized along the fibril axis (Fig. 8B). CD and secondary structure analysis indicate the predominance of antiparallel β-sheets in the fibrils of this study (Table S2; Table S3). Given the elongation process using dimers as the fundamental unit, it is concluded that the formed fibrils inevitably adopt a “double-stranded β-sheet” structure. In other words, the fibrils in solution are characterized by “double-stranded antiparallel β-sheets running parallel to the fibril axis.” This fiber-axis-parallel β-sheet structure is consistent with key features reported for native dragline silk (1, 4–9), indicating that our fibrils recapitulate major aspects of the native nanofibril architecture, at least in solution.

### Dehydration-Induced Transition and Polymorphism

Post-dehydration XRD patterns (Fig. 7) clearly indicate that this ordered fiber-axis-parallel β-structure is lost, with conversion to a cross-β state typical of amyloid fibrils, in which β-strands are oriented perpendicular to the fiber axis (Fig. 8C). If a cross-β structure had already formed in solution, the distance between N-termini would be drastically shorter than the periodic length estimated by AFM (approximately 30 nm)—specifically, 5.6 nm (calculated as 0.47 nm inter-strand spacing × 12 repeats). These two outcomes were obtained from the same fibrils dried under different boundary conditions, and we regard the comparison as informative rather than as a discrepancy to be explained away. In the AFM samples, fibrils are dried while adsorbed and fixed onto a mica substrate; interactions with the substrate physically restrict rearrangement of the molecular chains, and the ∼31.3 nm axial registry formed in solution is retained (Fig. 2 A and C). In contrast, the hydrogel used for XRD underwent macroscopic shrinkage and dried with far greater conformational freedom, and the axial architecture was lost in favour of cross-β (Fig. 7). The two observations rest on different observables — axial periodicity by AFM and crystalline architecture by XRD — so the comparison indicates rather than proves that the solution-state organization survives when chain mobility is restricted. It nevertheless identifies chain mobility during drying as the variable that must be controlled, and this is the basis of the two-stage design principle proposed below.

Why does such a drastic transition occur? We attribute the driving force to the alteration of the energy landscape associated with changes in hydration state. In solution, the intermolecular hydrogen-bond network presumably adopts a metastable state mediated by water molecules, maintaining an axial β-structure. However, when hydration water is removed by drying, the molecular chains are forced to rearrange in search of denser packing and more favorable hydrogen-bond networks. At this time, the system can be interpreted as falling into a “thermodynamic sink”—the cross-β structure, which is a deeper energetic minimum—as poly-Ala regions within molecules form strong hydrogen-bond networks and hydrophobic side chains pack between β-sheets.

Interestingly, a similar structural transition between the fiber-axis-parallel and cross-β architectures has been reported in the egg stalk silk of the lacewing *Chrysopa flava* (48). Geddes et al. found that the silk adopts a cross-β structure before stretching but converts to a fiber-axis-parallel β-sheet structure upon mechanical stretching after wetting. Although our system exhibits the reverse transition (fiber-axis-parallel → cross-β), both results suggest that silk proteins possess intrinsic structural polymorphism, allowing them to shuttle between at least two stable or metastable states: the “fiber-axis-parallel state” and the “cross-β state.”

Notably, we observed no reversible transition from cross-β to a fiber-axis-parallel structure upon mechanical stretching of the dried specimens until rupture. This is likely because the cross-β network derived from poly-Ala in our system is more robust than that in lacewing egg stalk silk, causing the dried specimens to break before intermolecular hydrogen bonds can rupture sufficiently.

Thus, the core challenge in artificial spinning hinges on how to suppress relaxation of the system to the thermodynamically most stable state (cross-β) during the inevitable drying process and instead fix it in the mechanically advantageous metastable state (the fiber-axis-parallel state). Consistently, our AFM observations show that immobilization on a substrate allows drying while fixing the solution-state structure, highlighting that “controlling molecular chain mobility” during the dehydration process is key to maintaining the native-like structure. In essence, controlling structural polymorphism not only during the self-assembly process itself but also during the subsequent dehydration and solidification process is central to the design of artificial spider silk comparable to native silk.

### Implications for *In Vivo* Spinning and Artificial Reconstruction

The pathway revealed in this study—fibrillization via a metastable dimer—provides insight into in vivo spinning mechanisms that extends beyond in vitro self-assembly. Two physical requirements underlie formation of the dimer, and it is these, rather than the particular solvent used, that have to be met. The first is that water be excluded from the contact surfaces, so that side chains can interdigitate at a solvent-excluded interface; in the preparation used here this is achieved by lowering the water activity of the solvent. The second is that main-chain hydrogen bonding remain suppressed while this occurs, so that association is arrested at the dimer rather than proceeding to disordered β-sheet stacking and higher-order oligomerization. Ethanol precipitation from concentrated GdmSCN satisfies both requirements by dehydration. Ethanol itself does not exist in vivo, and the pertinent question is therefore whether the same two requirements can be met by a different route.

They can be met without lowering the water activity of the solvent. Applying shear to the monomeric GU preparation, under conditions that otherwise yield no fibrils, confers fibrillogenic competence, and the extent of the reaction increases with the duration of the shear applied (Fig. S10). Flow within the spinning duct — both elongational and shear — has long been recognized as a driving force in spinning (23,32), and shear has been identified as a likely trigger for spidroin assembly (41). Access to the fibrillogenic state is therefore not confined to the route that lowers the water activity of the solvent.

Within the gland, several factors are expected to act in the same direction. Spidroin is stored at very high concentration in a hydrated gel, and liquid–liquid phase separation (LLPS) has been observed in vivo (33, 35, 49); within LLPS droplets the protein concentration is extremely high, bringing intermolecular distances close to the van der Waals contact distance and correspondingly reducing the hydration available to each molecule. Water is progressively extracted along the spinning duct (22). The N- and C-terminal domains, together with the pH and ion gradients, control the association state of spidroin (36–38,41), and would serve in vivo the role played by the denaturant in vitro, suppressing excessive main-chain aggregation while permitting specific side-chain contacts. In such an environment, adjacent poly-Ala regions would locally interdigitate to form a structure that excludes water molecules, similar to the core of the dimer observed here.

We should be explicit about what is and is not established. Within the system examined here, the preformed dimer is required: in its absence (GU), no fibrils are obtained at all. That an equivalent species acts as an intermediate in vivo is a proposal, not a demonstration. Testing it directly would require detection of the corresponding species within gland-derived material, which the present work does not attempt.

Furthermore, the fibril formation model obtained in this study provides a framework that reconciles the “micelle/LLPS hypothesis” and the “liquid crystal hypothesis,” which have traditionally been discussed as opposing views. Previously, spidroin was thought to form droplets via micelles or liquid–liquid phase separation (LLPS) at high concentrations, though the mechanistic details of their transformation into fibers remained elusive (33). On the other hand, liquid-crystalline behavior is observed in the spinning duct and is emphasized as a driving force for orientation (23, 32). The present results suggest the unification of both phenomena: droplets are not merely storage forms, but their interiors likely serve as “reaction chambers” where nuclei like metastable dimers are formed and grow into fibrils. The resulting nanofibrils (rigid rod-like particles) then behave as basic units (mesogens) for forming a liquid-crystalline phase, orienting along the fiber axis under flow.

Consistent with the proposed integrated model, Parent *et al.* reported the presence of fibril-like structures inside droplets within the gland (35), a finding that aligns well with our integrated model in which fibrils grow inside droplets and then form liquid crystals that align under flow. In other words, micelles/LLPS are responsible for the “supply of metastable dimers and the site of fibril growth,” whereas the liquid-crystalline phase provides the “orientation field of already formed nanofibrils,” allowing both to be repositioned as different aspects of a single continuous hierarchical process.

These findings provide a basis for molecular design principles for artificial spider silk. The core principle is to decouple self-assembly from dehydration and to optimize each independently. Metastable dimers and native-like fibrils with a fiber-axis-parallel β-structure must first be formed reliably under appropriate solvent and water-activity conditions; the subsequent drying and solidification then require physical intervention—mechanical constraint, rapid fixation, or orientation fixation by an external field—to prevent relaxation into the thermodynamically most stable cross-β structure. Spider silk reconstitution thus becomes a two-stage optimization problem. The picture of the metastable dimer and its dehydration-induced transition presented here provides molecular-scale principles for that problem, and may inform control strategies for amyloid-like structural polymorphism more generally.

## Materials and Methods

### Protein Preparation

Recombinant spidroin derived from the repetitive sequence of ADF3 (Araneus diadematus fibroin 3) was expressed in Escherichia coli and purified by Ni-affinity chromatography. To remove nucleic acids and induce the formation of metastable dimers, the purified protein was subjected to ethanol precipitation as detailed in SI Appendix.

### Sample Preparation

Two types of spidroin solutions with different solvation histories were prepared: “UU” (dissolved directly in 6 M urea) and “GU” (dissolved in 5 M guanidinium thiocyanate and dialyzed into 6 M urea). For solid-state analysis, oriented fibril samples were prepared by dialyzing UU solutions against 3 M urea and water to form hydrogels, which were subsequently dried with both ends mechanically fixed.

### Fibrillogenesis and Kinetic Assays

Fibrillogenesis was initiated by diluting stock solutions (in 6 M urea) twofold to a final concentration of 3 M urea. The reaction kinetics were monitored by Thioflavin T (ThT) fluorescence.

### Structural Characterization

Solution structures of the precursors and fibrils were analyzed by small-angle X-ray scattering (SAXS) at the Photon Factory (KEK) and circular dichroism (CD) spectroscopy. Fibril morphologies were characterized by atomic force microscopy (AFM). Real-time observation of fibril elongation was performed using high-speed AFM (HS-AFM). The internal structure of dried fibrils was analyzed by wide-angle X-ray diffraction (XRD). To isolate anisotropic crystalline reflections, difference profiles were calculated by subtracting equatorial profiles from meridional profiles to remove isotropic amorphous scattering. Detailed protocols for all experiments are provided in SI Appendix.

## Data Availability

The data underlying this article are available in the article and in its online supplementary material. Additional data underlying this article will be shared on reasonable request to the corresponding author.

## Acknowledgments

We are grateful to Takuya Sawai for his preliminary work that laid the foundation for this study. Synchrotron radiation experiments were performed at the Photon Factory of the High Energy Accelerator Research Organization (KEK) under Proposal Nos. 2016G077, 2018G119, 2020G064, and 2023G053. This work was supported by the Impulsing Paradigm Change through Disruptive Technologies (ImPACT) Program of the Council for Science, Technology and Innovation (Cabinet Office, Government of Japan); Japan Science and Technology Agency (JST) A-STEP Grant Number JPMJTR20UV (to H.K.); and JSPS KAKENHI Grant Number JP23H02448 and 22H04984 (to H.K.).

## Use of AI

During preparation of this revision, the authors used Anthropic’s Claude (Opus 4.8, 5) to assist with English language editing, with drafting and redrafting of text in the revised manuscript and the point-by-point response, and with checking internal consistency across the revised files. The tool was not used to generate, analyze, or interpret experimental data, nor to produce any figure or image. The authors reviewed and verified all content and take full responsibility for the published work.

## Author Contributions

Kamikubo and H. Kajimoto designed research; H. Kamikubo and H. Kajimoto developed the methodology; H. Kajimoto, K.Y., K.S.C., K.H., Y.O., R.A., Y.N., K.K., M.B., Y.Y., S.T.F., T.U., and T.K.S. performed research; H. Kamikubo, H. Kajimoto, and K.Y. analyzed data; H. Kamikubo and H. Kajimoto wrote the paper; and H. Kamikubo, H. Kajimoto, K.Y., Y.Y., and S.T.F. reviewed and edited the manuscript.

## References

1. J. M. Gosline, M. E. DeMont, M. W. Denny, The structure and properties of spider silk. Endeavour 10, 37–43 (1986).

2. J. E. Garb, et al., The transcriptome of Darwin’s bark spider silk glands predicts proteins contributing to dragline silk toughness. *Commun*. Biol. 2, 275 (2019).

3. R. V. Lewis, Spider silk: the unraveling of a mystery. Acc. Chem. Res. 25, 392–398 (1992).

4. S. Sampath, et al., X-ray diffraction study of nanocrystalline and amorphous structure within major and minor ampullate dragline spider silks. Soft Matter 8, 6713–6722 (2012).

5. J. O. Warwicker, Comparative studies of fibroins: II. The crystal structures of various fibroins. J. Mol. Biol. 2, 350–362 (1960).

6. A. D. Parkhe, S. K. Seeley, K. Gardner, L. Thompson, R. V. Lewis, Structural studies of spider silk proteins in the fiber. J. Mol. Recognit. 10, 1–6 (1997).

7. A. Bram, C. I. Brändén, C. Craig, I. Snigireva, C. Riekel, X-ray diffraction from single fibres of spider silk. J. Appl. Crystallogr. 30, 390–392 (1997).

8. Z. Dong, R. V. Lewis, C. R. Middaugh, Molecular mechanism of spider silk elasticity. Arch. Biochem. Biophys. 284, 53–57 (1991).

9. A. Simmons, E. Ray, L. W. Jelinski, Solid-state 13C NMR of Nephila clavipes dragline silk establishes structure and identity of crystalline regions. Macromolecules 27, 5235–5237 (1994).

10. S. Xiao, S. Xiao, F. Gräter, Dissecting the structural determinants for the difference in mechanical stability of silk and amyloid beta-sheet stacks. Phys. Chem. Chem. Phys. 15, 8765–8771 (2013).

11. E. Oroudjev, et al., Segmented nanofibers of spider dragline silk: atomic force microscopy and single-molecule force spectroscopy. Proc. Natl. Acad. Sci. U. S. A. 99 Suppl 2, 6460–6465 (2002).

12. V. G. Bogush, et al., A novel model system for design of biomaterials based on recombinant analogs of spider silk proteins. J. Neuroimmune Pharmacol. 4, 17–27 (2009).

13. T. K. Tenchurin, et al., Effect of recombinant spidroins self-assembly on rheological behavior of their dispersions and structure of electrospun nanofibrous materials. Polymers (Basel) 15, 3001 (2023).

14. C.-F. Hu, C.-Y. Gan, Y.-J. Zhu, X.-X. Xia, Z.-G. Qian, Modulating polyalanine motifs of synthetic spidroin for controllable preassembly and strong fiber formation. ACS Biomater. Sci. Eng. 10, 2925–2934 (2024).

15. D. Huemmerich, et al., Novel assembly properties of recombinant spider dragline silk proteins. Curr. Biol. 14, 2070–2074 (2004).

16. M. Humenik, M. Magdeburg, T. Scheibel, Influence of repeat numbers on self-assembly rates of repetitive recombinant spider silk proteins. J. Struct. Biol. 186, 431–437 (2014).

17. M. Humenik, A. M. Smith, S. Arndt, T. Scheibel, Ion and seed dependent fibril assembly of a spidroin core domain. J. Struct. Biol. 191, 130–138 (2015).

18. U. Slotta, et al., Spider silk and amyloid fibrils: a structural comparison. Macromol. Biosci. 7, 183–188 (2007).

19. C. W. P. Foo, et al., Role of pH and charge on silk protein assembly in insects and spiders. Appl. Phys. A Mater. Sci. Process. 82, 223–233 (2006).

20. M. Andersson, et al., Carbonic anhydrase generates CO2 and H+ that drive spider silk formation via opposite effects on the terminal domains. PLOS Biol. 12, e1001921 (2014).

21. D. P. Knight, F. Vollrath, Changes in element composition along the spinning duct in a Nephila spider. Naturwissenschaften 88, 179–182 (2001).

22. E. K. Tillinghast, S. F. Chase, M. A. Townley, Water extraction by the major ampullate duct during silk formation in the spider, Argiope aurantia Lucas. J. Insect Physiol. 30, 591–596 (1984).

23. D. P. Knight, F. Vollrath, Liquid crystals and flow elongation in a spider’s silk production line. Proc. R. Soc. Lond. B Biol. Sci. 266, 519–523 (1999).

24. S. Sonavane, P. Westermark, A. Rising, L. Holm, Regionalization of cell types in silk glands of Larinioides sclopetarius suggest that spider silk fibers are complex layered structures. Sci. Rep. 13, 22273 (2023).

25. J. M. Kenney, D. Knight, M. J. Wise, F. Vollrath, Amyloidogenic nature of spider silk. Eur. J. Biochem. 269, 4159–4163 (2002).

26. J. L. Yarger, B. R. Cherry, A. van der Vaart, Uncovering the structure–function relationship in spider silk. Nat. Rev. Mater. 3, 18008 (2018).

27. Q. Wang, H. C. Schniepp, Nanofibrils as building blocks of silk fibers: critical review of the experimental evidence. JOM 71, 1248–1263 (2019).

28. D. Perera, et al., Natural spider silk nanofibrils produced by assembling molecules or disassembling fibers. Acta Biomater. 168, 323–332 (2023).

29. Q. Wang, H. C. Schniepp, Strength of recluse spider’s silk originates from nanofibrils. ACS Macro Lett. 7, 1364–1370 (2018).

30. Q. Wang, et al., Observations of 3 nm silk nanofibrils exfoliated from natural silkworm silk fibers. ACS Mater. Lett. 2, 153–160 (2020).

31. A. A. Walker, C. Holland, T. D. Sutherland, More than one way to spin a crystallite: multiple trajectories through liquid crystallinity to solid silk. Proc. Biol. Sci. 282, 20150259 (2015).

32. F. Vollrath, D. P. Knight, Liquid crystalline spinning of spider silk. Nature 410, 541–548 (2001).

33. A. D. Malay, et al., Spider silk self-assembly via modular liquid-liquid phase separation and nanofibrillation. Sci. Adv. 6, eabb6030 (2020).

34. A. D. Malay, H. C. Craig, J. Chen, N. A. Oktaviani, K. Numata, Complexity of spider dragline silk. Biomacromolecules 23, 1827–1840 (2022).

35. L. R. Parent, et al., Hierarchical spidroin micellar nanoparticles as the fundamental precursors of spider silks. Proc. Natl. Acad. Sci. U. S. A. 115, 11507–11512 (2018).

36. M. Landreh, et al., A pH-dependent dimer lock in spider silk protein. J. Mol. Biol. 404, 328–336 (2010).

37. G. Askarieh, et al., Self-assembly of spider silk proteins is controlled by a pH-sensitive relay. Nature 465, 236–238 (2010).

38. N. Kronqvist, et al., Sequential pH-driven dimerization and stabilization of the N-terminal domain enables rapid spider silk formation. Nat. Commun. 5, 3254 (2014).

39. J. Bauer, et al., Acidic residues control the dimerization of the N-terminal domain of black widow spiders’ major ampullate spidroin 1. Sci. Rep. 6, 34442 (2016)

40. M. Sarr, et al., The dimerization mechanism of the N-terminal domain of spider silk proteins is conserved despite extensive sequence divergence. J. Biol. Chem. 298, 101913 (2022)

41. F. Hagn, et al., A conserved spider silk domain acts as a molecular switch that controls fibre assembly. Nature 465, 239–242 (2010)

42. H. R. Johnson, et al., Distinct dimerization mechanisms in silkworm and spider silk proteins revealed by mass photometry. J. Phys. Chem. Lett. 16, 11451–11457 (2025).

43. M. Allan, L. J. Mauer, Dataset of water activity measurements of alcohol: water solutions using a tunable diode laser. Data Brief 12, 364–369 (2017).

44. E. Tsunekawa, et al., X-ray and electron diffraction observations of steric zipper interactions in metal-induced peptide cross-β nanostructures. J. Am. Chem. Soc. 145, 16160–16165 (2023).

45. T. Asakura, et al., Two different packing arrangements of antiparallel polyalanine. Angew. Chem. Int. Ed Engl. 51, 1212–1215 (2012).

46. T. Asakura, Y. Tasei, A. Aoki, A. Nishimura, Mixture of rectangular and staggered packing arrangements of polyalanine region in spider dragline silk in dry and hydrated states as revealed by 13C NMR and X-ray diffraction. Macromolecules 51, 1058–1068 (2018).

47. M. El Guendouzi, A. Benbiyi, Thermodynamic properties of binary aqueous solutions of orthophosphate salts, sodium, potassium and ammonium at T=298.15K. Fluid Phase Equilib. 369, 68–85 (2014).

48. A. J. Geddes, K. D. Parker, E. D. Atkins, E. Beighton, “Cross-beta” conformation in proteins. J. Mol. Biol. 32, 343–358 (1968).

49. T.-Y. Lin, et al., Liquid crystalline granules align in a hierarchical structure to produce spider dragline microfibrils. Biomacromolecules 18, 1350–1355 (2017).

