## Supplementary for "Metastable Dimeric Intermediate Drives Native-like Spidroin Fibrillogenesis, while Dehydration Promotes Cross-β Transition"

**This PDF file includes:**

Supporting text
Figures S1 to S10
Tables S1 to S4
References
Legends for Movies S1

**Other supporting materials for this manuscript include the following:**

Movies S1

### Supporting Information Text

### Materials and Methods

#### 1. Protein Preparation

Recombinant spidroin derived from the repetitive sequence of ADF3 (*Araneus diadematus* fibroin 3), a major dragline silk protein of *Araneus diadematus*, was used in this study. The amino acid sequence and molecular weight of the construct are provided below. The protein was expressed in *Escherichia coli* and purified by Ni-affinity chromatography using an N-terminal His tag.

**Amino acid sequence of the spidroin**
Molecular Weight: 47.5 kDa
Number of Residues: 542

```
MHHHHHHHHHHSSGSSLEVLFQGPARAGSGQQGPGQQGPGQQGPGQQGPY
GPGASAAAAAAGGYGPGSGQQGPSQQGPGQQGPGGQGPYGPGASAAAAAA
GGYGPGSGQQGPGGQGPYGPGSSAAAAAAGGNGPGSGQQGAGQQGPGQQG
PGASAAAAAAGGYGPGSGQQGPGQQGPGGQGPYGPGASAAAAAAGGYGPG
SGQGPGQQGPGGQGPYGPGASAAAAAAGGYGPGSGQQGPGQQGPGQQGPG
GQGPYGPGASAAAAAAGGYGPGYGQQGPGQQGPGGQGPYGPGASAASAAS
GGYGPGSGQQGPGQQGPGGQGPYGPGASAAAAAAGGYGPGSGQQGPGQQG
PGQQGPGQQGPGGQGPYGPGASAAAAAAGGYGPGSGQQGPGQQGPGQQGP
GQQGPGQQGPGQQGPGQQGPGQQGPGQQGPGGQGAYGPGASAAAGAAGGY
GPGSGQQGPGQQGPGQQGPGQQGPGQQGPGQQGPGQQGPGQQGPYGPGAS
AAAAAAGGYGPGSGQQGPGQQGPGQQGPGGQGPYGPGAASAA
```

To remove nucleic acids from the obtained spidroin powder, ethanol precipitation was performed. First, the lyophilized powder was dissolved in 5 M guanidinium thiocyanate (GdmSCN) solution (10 mM Tris-HCl, 5 mM DTT, pH 7.0) to a concentration of 100 mg/mL. Ethanol was added dropwise to this solution to achieve stepwise concentrations of 50, 60, and 75 vol%, with centrifugation (15,000 × *g*, 20°C, 10 min) performed at each step. Sodium dodecyl sulfate–polyacrylamide gel electrophoresis (SDS-PAGE) confirmed that spidroin was recovered mainly in the precipitate fraction upon addition of 75 vol% ethanol. The precipitate was washed five times with ultrapure water to remove salts, resuspended in ultrapure water, frozen in liquid nitrogen, and lyophilized. The resulting "purified powder" was used for subsequent experiments.

#### 2. Sample Preparation

**Urea Solubility Test.** To evaluate the solubility of the purified powder, the powder was added to 0–8 M urea solutions (10 mM Tris-HCl, 5 mM DTT, pH 7.0) at 25 mg/mL. After vortexing and standing overnight, undissolved material was removed by centrifugation (20,000 × *g*, 20°C, 20 min). The protein concentration in the supernatant was determined from the absorbance at 280 nm using the molar extinction coefficient calculated from the amino acid sequence (S1).

**Preparation of UU and GU Solutions.** Two types of spidroin solutions with different solvation histories (UU and GU) were prepared in this study.

**UU (urea–urea)**: The purified powder was added directly to 6 M urea solution (10 mM Tris-HCl, 5 mM DTT, pH 7.0) at 25 mg/mL, vortexed, and allowed to dissolve overnight at room temperature. The solution was then dialyzed (MWCO 8,000) at room temperature against the same 6 M urea solution used for GU dialysis to exchange the solvent. After removal of trace amounts of undissolved material by centrifugation (20,000 × *g*, 20°C, 20 min), the supernatant was sonicated for 1 min and used as the UU stock solution.

**GU (Guanidine–urea)**: The purified powder was dissolved in 5 M GdmSCN solution (10 mM Tris-HCl, 5 mM DTT, pH 7.0) to fully denature the structure, and then dialyzed (MWCO 8,000) at room temperature against 6 M urea solution (same buffer conditions) to exchange the solvent. The resulting solution was used as the GU stock solution.

**Preparation of Oriented Fibril Hydrogels.** For X-ray diffraction (XRD) measurements, the UU stock solution (1.5 mL) was sealed in a dialysis tube (MWCO 8,000) with clips to increase internal pressure and dialyzed against 3 M urea solution (10 mM Tris-HCl, 5 mM DTT, pH 7.0) at room temperature for 5 days to induce fibrillogenesis. Dialysis was then continued against ultrapure water for 7 days to obtain a desalted hydrogel. The hydrogel was removed from the tube, mounted with both ends fixed, and dried at room temperature, thereby producing a uniaxially oriented solid sample by utilizing the shrinkage stress generated during drying.

### 3. Fibrillogenesis Assay

**Thioflavin T (ThT) Kinetics.** The time course of the fibrillogenesis reaction was monitored by ThT fluorescence measurements. Measurements were performed using a plate reader (Synergy HTX, Agilent Technologies) with an excitation wavelength of 440 nm, an emission wavelength of 500 nm, and a temperature of 27 °C, acquiring data at 15-min intervals. The reaction was initiated by twofold dilution of the UU and GU stock solutions with a dilution buffer containing ThT (10 mM Tris-HCl, 5 mM DTT, pH 7.0; final conditions: 3 M urea, 4 μM ThT). The fluorescence intensities obtained were background-corrected by subtracting the values measured for the corresponding buffer alone.

### 4. Structural Characterization

**Atomic Force Microscopy (AFM).** Morphological observation was performed by tapping-mode AFM in air using an SPM-9700HT (Shimadzu). An OMCL-AC240TS-R3 cantilever (Olympus; spring constant 1.7 N/m, resonant frequency 70 kHz) was used. Samples were prepared by depositing a spidroin solution (0.005–1 mg/mL) onto a mica substrate, allowing adsorption for 30 s, rinsing with ultrapure water, and drying at room temperature. For gold nanoparticle labeling experiments to confirm fibril orientation, a solution of Ni-NTA-functionalized gold nanoparticles (5 nm diameter, Nanoprobes) was deposited onto mica with adsorbed spidroin, incubated for 10 min, rinsed and dried.

**Small-Angle X-ray Scattering (SAXS).** Static solution structural analysis (UU vs. GU) was performed at 25 °C at beamline BL-10C of the Photon Factory, High Energy Accelerator Research Organization (KEK) (wavelength 1 Å, camera length 2 m). Time-resolved measurements (tracking fibril elongation) were performed using a laboratory X-ray source (MicroMAX-007HF, Rigaku) equipped with a Pilatus 200K detector. The obtained two-dimensional scattering images were converted into one-dimensional intensity profiles, $I(Q)$, by circular averaging. Solvent-subtracted profiles were used for Guinier analysis and for constructing Kratky plots.

**Circular Dichroism (CD).** Secondary structure analysis was performed using a circular dichroism spectropolarimeter (J-725, JASCO). Measurements were carried out using quartz cells with a path length of 0.01 cm (for high-concentration samples) or 0.1 cm (for low-concentration samples) over a wavelength range of 200–250 nm, with a scan rate of 20 nm/min at 20 °C. After subtracting the buffer contribution from the obtained spectra, secondary structure contents were estimated using the CDPro software package (algorithm: CONTINLL, reference set: SDP48) (S2) and the BeStSel server (S3).

**X-ray Diffraction (XRD).** Wide-angle X-ray diffraction measurements of dried hydrogels were performed at room temperature using a MicroMAX-007HF (Rigaku) X-ray generator and an R-AXIS

VI imaging plate detector. The X-ray wavelength was Cu Kα (1.54 Å), and the exposure time was 900 s. One-dimensional profiles along the meridional (fiber-axis) and equatorial directions were extracted from the obtained two-dimensional diffraction images, and background correction was performed using air scattering. To isolate the anisotropic crystalline reflections from the isotropic amorphous halo, a difference profile was calculated by subtracting the equatorial profile from the meridional profile (Fig. S9). For analysis of the cross-β structure, peak deconvolution using Gaussian functions was performed to determine the *d*-spacings of the major reflections.

**High Speed AFM (HS-AFM).** HS-AFM observations were conducted in tapping mode using a laboratory-built instrument. An Olympus BL-AC7 cantilever, with a nominal spring constant of approximately 0.2 N/m, was utilized. The tips were prepared by depositing amorphous carbon onto the apex of the cantilever using focused electron beam exposure, followed by sharpening via plasma etching in argon gas. For *in situ* observation of fibril growth, 0.5 mg/mL precursor protein solution was first applied onto a freshly cleaved bare mica substrate to allow for protein adsorption. The substrate was then gently rinsed with the observation buffer (10 mM Tris-HCl, 5 mM DTT) to remove unbound protein, and the high-speed AFM imaging was initiated in this buffer. To trigger and record the fibril elongation process in real time, 20 μL of the precursor solution, prepared in 6 M urea, was injected into the observation chamber, resulting in a final urea concentration of approximately 1.7 M (total volume, ~70 μL). The imaging speed and scan area for each experiment are specified in the corresponding movie legends.

**Figures**

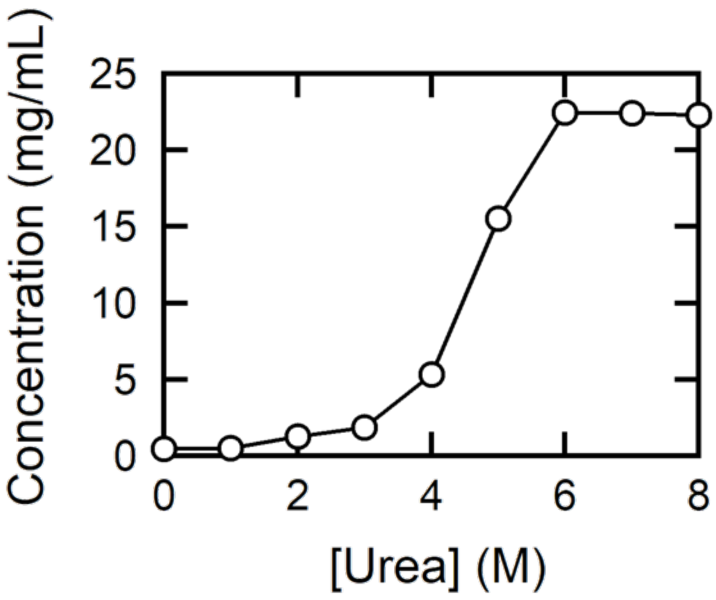


**Fig. S1. Solubility of purified spidroin powder at different urea concentrations.** High solubility (>20 mg/mL) was observed at urea concentrations of 6 M or higher, whereas aggregation increased sharply at lower concentrations. In 0 M urea (ultrapure water), the solubility decreased to approximately 0.5 mg/mL (the midpoint of the solubility curve was approximately 4.5 M).

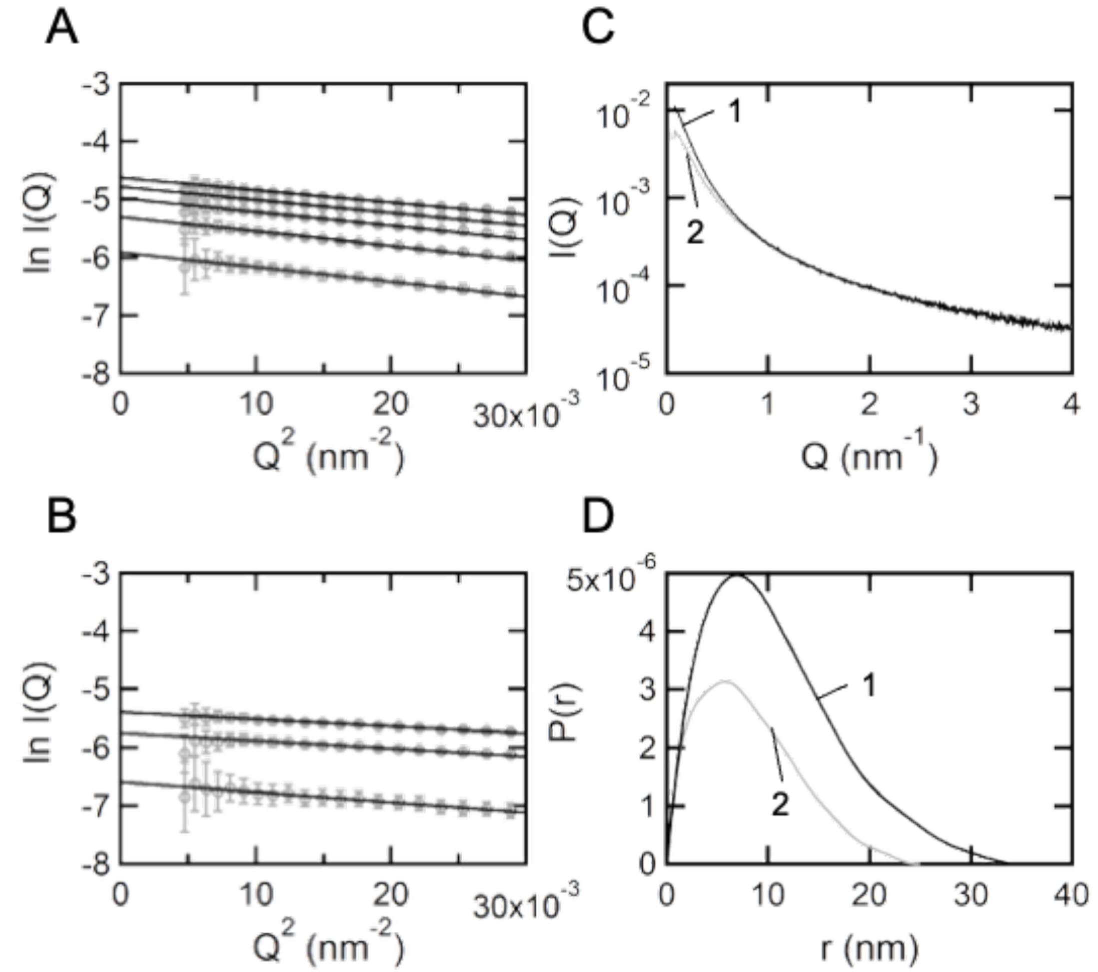


**Fig. S2. SAXS analysis of UU and GU in 6 M urea.** (A and B) Guinier plots of UU (A) and GU (B). Linear regression was performed in the range of $0.0021 < Q^2 < 0.00814$ nm$^{-2}$ to satisfy the condition $QRg < 1.3$. (C) Solvent-subtracted scattering profiles *I(Q)* for UU (Line 1) and GU (Line 2), shown on logarithmic axes. These are the primary data from which the Guinier plots in (A) and (B) and the Kratky plots in Fig. 3C were derived. (D) Distance distribution functions *P(r)* computed from the profiles in (C). The maximum dimensions $D_{max}$ were 35 nm for UU and 25 nm for GU. *P(r)* provides a comparison of relative molecular mass that is independent of the Guinier analysis, since the Guinier approximation uses only the low-angle region whereas *P(r)* is computed over a wide angular range. Normalized to concentration, the UU:GU ratio obtained from P(r) was 1.96, compared with 1.90 from the Guinier analysis (Table 1); both indicate that UU is a dimer and GU a monomer. UU, urea–urea; GU, guanidine–urea; *P(r)*, distance distribution function; $D_{max}$, maximum dimension.

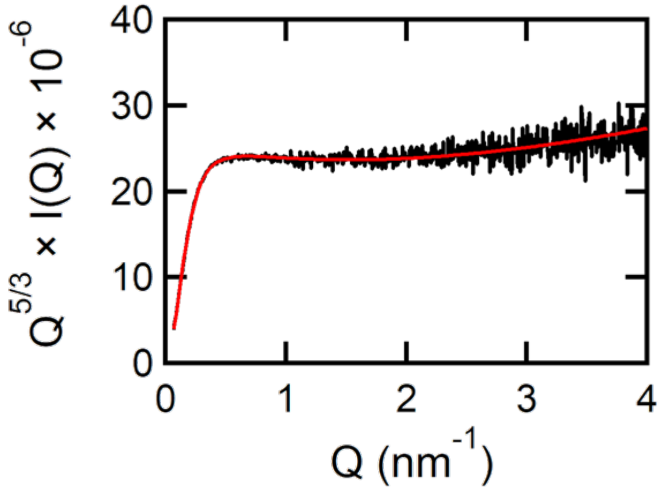


**Fig. S3. Fitting of the polymer excluded-volume model to the modified Kratky plot of GU.** The black line represents the experimental data, and the red line indicates the regression curve based on the polymer model with excluded volume (see below). Fitting was performed without fixing any parameters. As a result, the radius of gyration $R_g$ was calculated to be 6.2 +/− 0.01 nm, which is consistent with the value obtained from the Guinier analysis. The excluded-volume parameter $\nu$ was estimated to be 0.549 +/− 0.001, and the mass fractal dimension $m$ (the reciprocal of $\nu$) was calculated to be 1.8. These results quantitatively indicate that GU behaves as a typical random coil expanded by excluded-volume effects. GU, guanidine–urea.

**Excluded volume polymer model.** The form factor $P(Q)$ of a polymer with excluded volume can be expressed using the excluded-volume parameter $\nu$ as follows (S4).

$$P(Q) = \frac{1}{\nu U^{1/2\nu}} \gamma\left(\frac{1}{2\nu}, U\right) - \frac{1}{\nu U^{1/\nu}} \gamma\left(\frac{1}{\nu}, U\right)$$

Here, $\gamma$ denotes the incomplete gamma function. $U = Q^2 R_g^2 (2\nu + 1)(2\nu + 2)/6$, and $R_g$ represents the radius of gyration. The mass fractal dimension $m$ can be expressed in terms of $\nu$ as $m = 1/\nu$. In practice, the following equation was used for fitting:

$$P(Q) = Scale \cdot P(Q) + Background$$

**As a reference for interpreting the Kratky plots in Fig. 3C:** a compact, globular particle gives a pronounced peak, whereas a random coil gives a plateau extending to wide angles. Star polymers and dendrimers, in which several flexible arms emanate from a compact center, give a superposition of the two — a low-angle peak arising from the core superimposed on the plateau arising from the arms — and the relative prominence of the low-angle peak increases with the volume fraction occupied by the compact center (S4, S5). GU corresponds to the random-coil limit, as quantified above. UU shows the same wide-angle plateau as GU but with an additional broad peak near Q ~ 0.32 nm$^{-1}$ (Fig. 3C), and therefore corresponds to the intermediate case characteristic of a branched architecture of low functionality — a star polymer with few arms, or a low-generation dendrimer — in which a compact region of limited volume fraction is surrounded by chains whose statistics are indistinguishable from those of the monomeric GU. We note the limits of this description: it specifies the presence and approximate size of a compact region and the statistics of the remaining chain, but not the conformation of the residues within either region, nor which residues occupy the core.

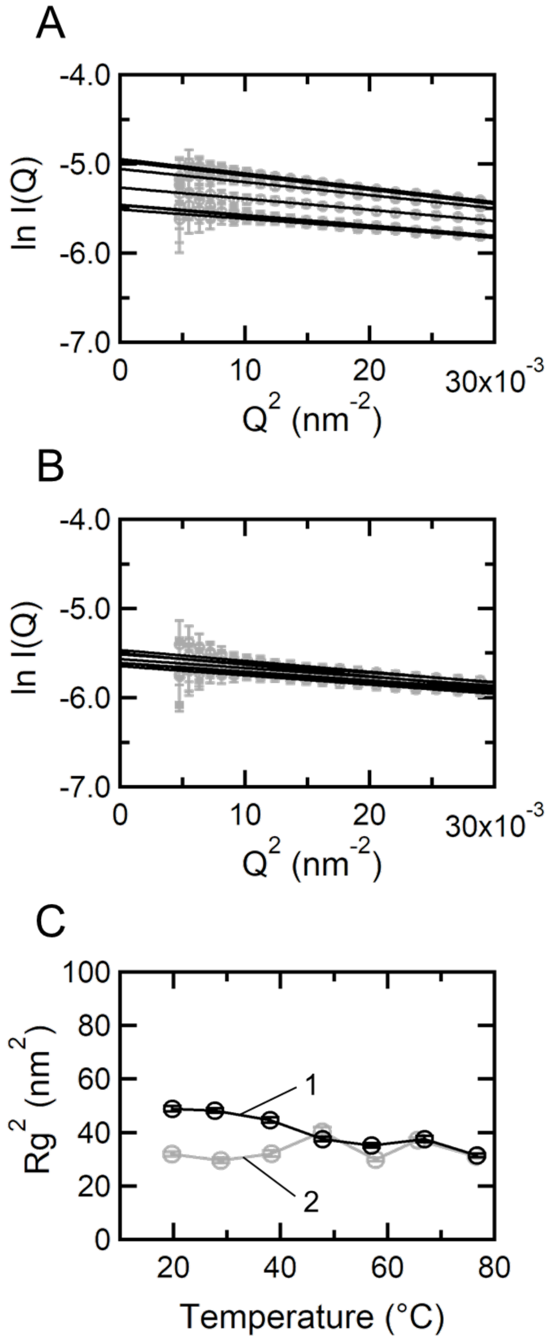


**Fig. S4. SAXS measurements of UU during heating and cooling processes in 6 M urea.** (*A* and *B*) Guinier plots during the heating (*A*) and cooling (*B*) processes. The regression range was set to 0.00345 < $Q^2$ < 0.00814 nm$^{-2}$ to satisfy the condition $Q\ R_g < 1.3$. (*C*) Changes in the $R_g$ obtained from Guinier analysis during heating (Line 1) and cooling (Line 2). Similar to the behavior of *I*(0)/*Conc.* (Fig. 4), $R_g$ decreased cooperatively between 40 °C and 60 °C during heating and did not return to the original value upon cooling. SAXS, small-angle X-ray scattering; UU, urea–urea; $R_g$, radius of gyration.

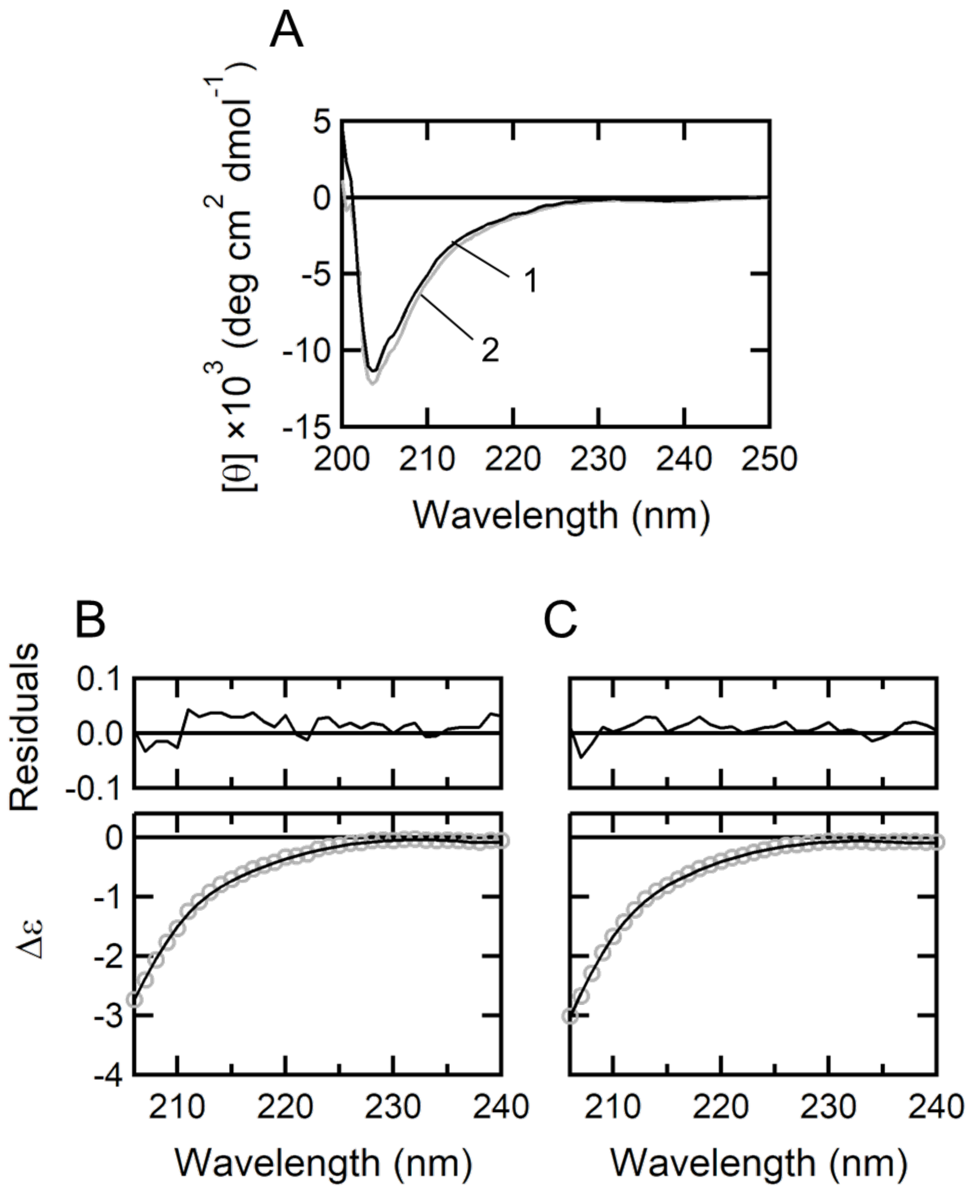


**Fig. S5. CD spectra and secondary structure analysis of UU and GU in 6 M urea.** (*A*) Comparison of CD spectra for UU (Line 1) and GU (Line 2) measured in 6 M urea. The two spectra are in good agreement, indicating that both samples are predominantly unstructured. (*B* and *C*) CD spectral analysis of UU (*B*) and GU (*C*) under 6 M urea conditions. Spectra were analyzed using the CONTINLL algorithm implemented in CDPro. Experimental data are shown as markers, and the black lines indicate the fitted curves (analysis range: 206–240 nm), with residuals shown above. The SDP48 reference dataset containing denatured proteins was used. CD, circular dichroism; UU, urea–urea; GU, guanidine–urea.

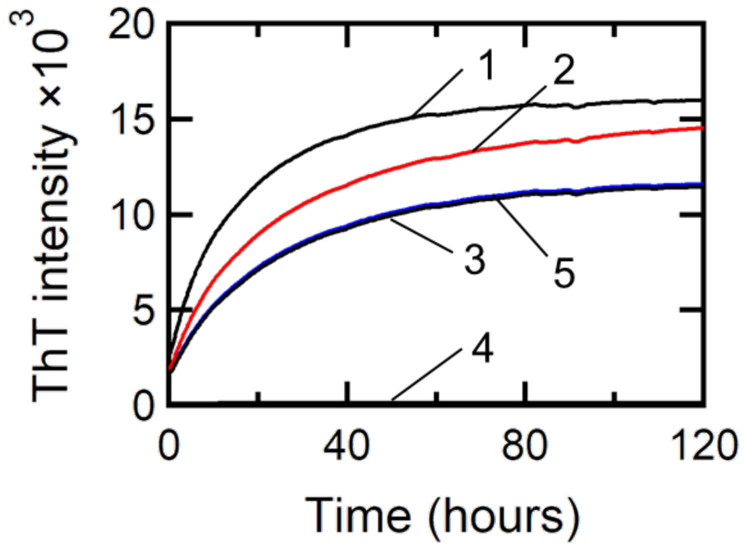


**Fig. S6. Time course of ThT fluorescence intensity for UU and GU mixtures in 3 M Urea**. In addition to a mixed system containing 10 mg/mL UU and 5 mg/mL GU (final concentrations; Line 2), the following samples were prepared for comparison: 15 mg/mL UU alone (Line 1), 10 mg/mL UU alone (Line 3), and 5 mg/mL GU alone (Line 4). Measurements were initiated immediately after twofold dilution with Tris buffer. If the dimers do not function as nuclei, the reaction curve of the mixture (Line 2) is expected to coincide with the arithmetic sum of the 10 mg/mL UU alone and 5 mg/mL GU alone systems (Line 5). Conversely, if the dimers function as nuclei and efficiently incorporate monomers into fibrils, the final fluorescence intensity of the mixture (Line 2) is expected to approach that of the 15 mg/mL UU alone system (Line 1). In practice, the fluorescence intensity curve of the mixture (Line 2) exhibited behavior intermediate between the 15 mg/mL UU curve (Line 1) and the sum of the 10 mg/mL UU and 5 mg/mL GU systems (Line 5). This result suggests that while UU functions as a nucleus, it does not efficiently incorporate monomers into fibrils, indicating that the conversion efficiency is limited. ThT, thioflavin T; UU, urea–urea; GU, guanidine–urea.

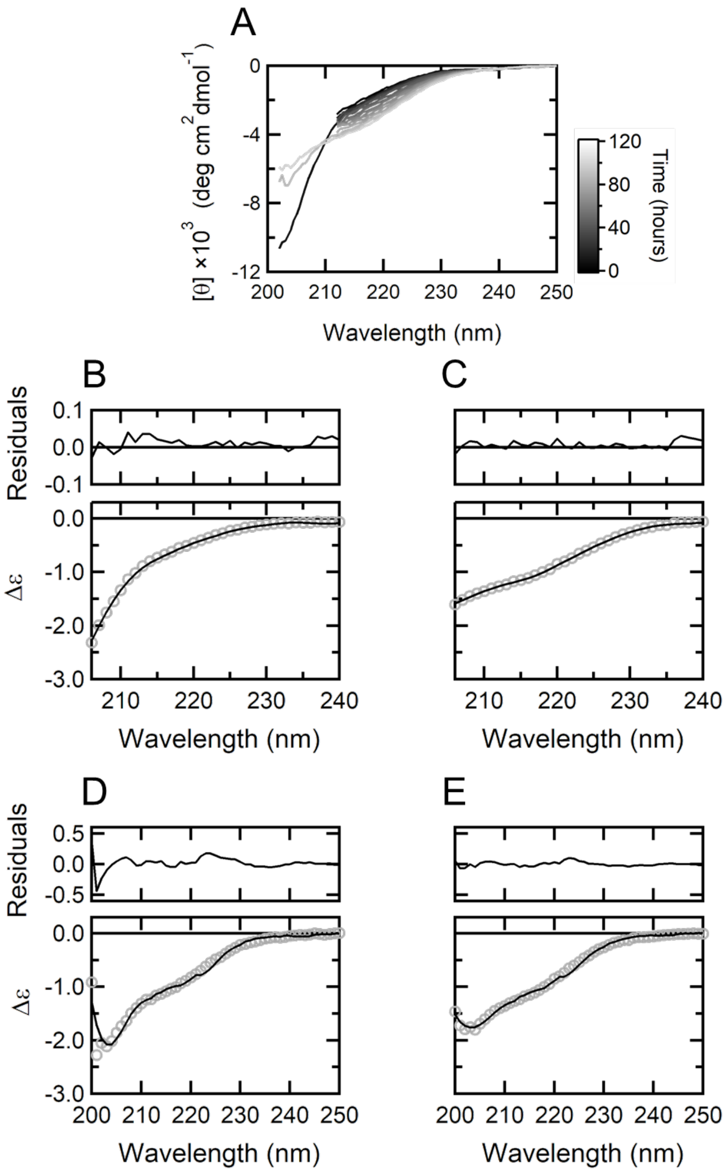


**Fig. S7. Time-resolved CD during fibril elongation.** (*A*) Time evolution of CD spectra from immediately after diluting UU to 3 M urea up to 120 h. The negative ellipticity around 210–230 nm increased over time, suggesting the formation of β-sheet structures accompanying fibril elongation. (*B* and *C*) CDPro analysis (CONTINLL, SDP48) of spectra for UU immediately after dilution into 3 M urea (*B*) and after 120 h (*C*) (analysis range: 206–240 nm). (*D* and *E*) BeStSel analysis of spectra immediately after dilution (*D*) and after 120 h (*E*) (analysis range: 200–250 nm). In all plots, experimental data are shown as markers, and the black lines indicate the fitted curves, with residuals shown above. Secondary structure contents estimated using BeStSel are summarized in Table S3. The results from both methods are generally consistent, suggesting that antiparallel β-sheets are formed as the major structure during fibril elongation. CD, circular dichroism; UU, urea–urea.

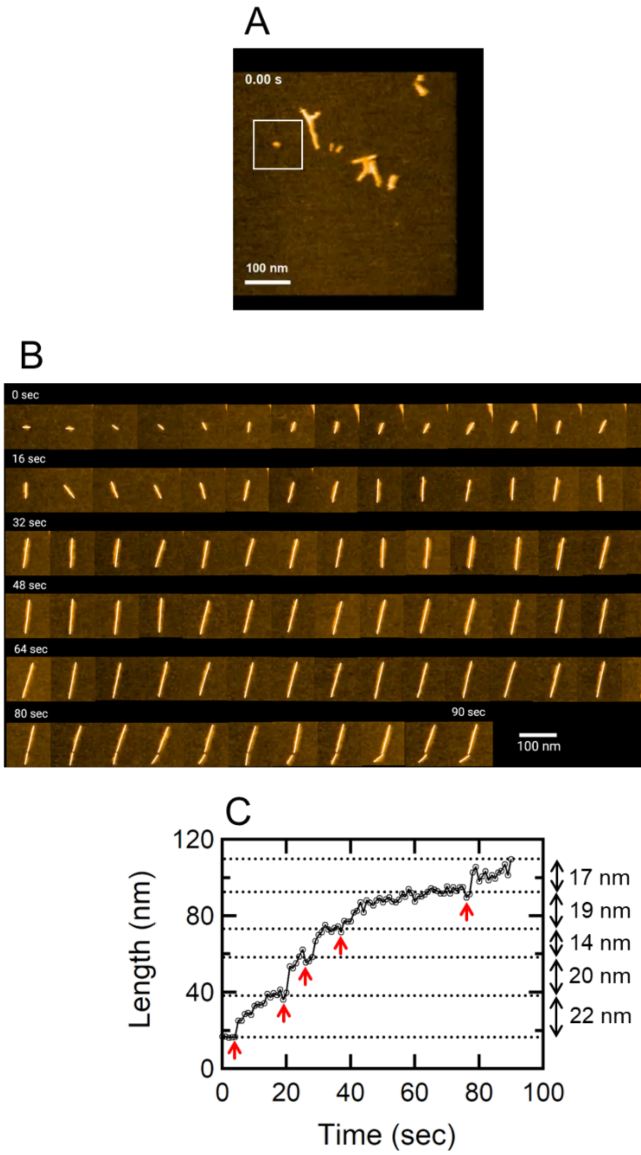


**Fig. S8. HS-AFM analysis of the fibril elongation process (Movie S1).** (*A*) Field of view in Movie S1. The white square indicates the initial position of the cropped area shown in (*B*). (*B*) Time-lapse images of fibril elongation (0–90 s). (*C*) Time course of fibril length. Red arrows indicate points where the length increased abruptly. Following these abrupt steps, a gradual increase in length was observed. This two-stage growth process suggests that the added nanoparticles (dimers) do not align immediately upon attachment; instead, they first loosely associate via the random-coil regions surrounding the core to form an "encounter complex." Subsequent structural rearrangement of the poly-Ala segments within the random coils likely drives β-sheet formation, leading to the maturation of the stable fibril structure. Notably, the increment per growth step observed by HS-AFM was smaller than the periodicity characteristic of fully matured fibrils (~31.3 nm; Fig. 2). We suggest that this difference is attributable to the degree of fibril maturation; whereas Fig. 2 captures the state after sufficient elongation (60 h), HS-AFM visualizes the nascent state within minutes of formation. Crucially, even at this immature stage, the step size is significantly larger than the theoretical length expected for a cross-β structure (~5.6 nm, see discussion). HS-AFM, high-speed atomic force microscopy.

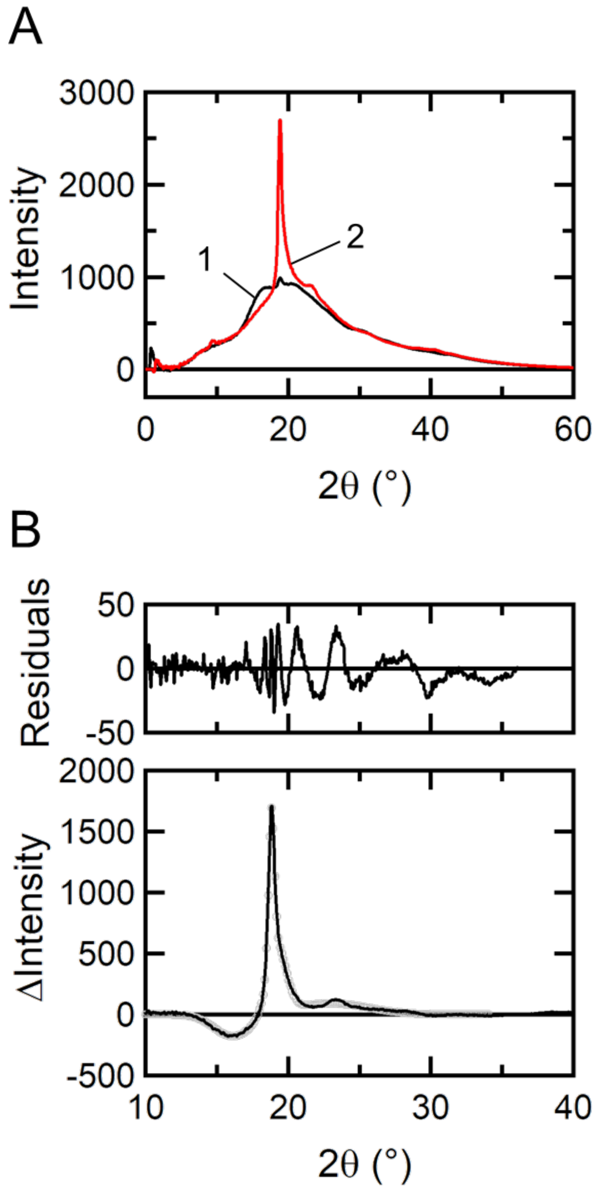


**Fig. S9. XRD of the stretched and dried hydrogel**. (*A*) One-dimensional profiles obtained by sector averaging the two-dimensional diffraction pattern (Fig. 7C) along the equatorial (Line 1) and meridional (Line 2) directions. (*B*) Difference profile (markers) obtained by subtracting the equatorial profile (Line 1) from the meridional profile (Line 2). This subtraction removes broad isotropic scattering derived from amorphous regions, clarifying the contribution of crystalline regions. The black line represents the fitting result using a four-component Gaussian function, and residuals are shown above. This analysis identified peaks characteristic of cross-β structures (see Table S4). XRD, X-ray diffraction.

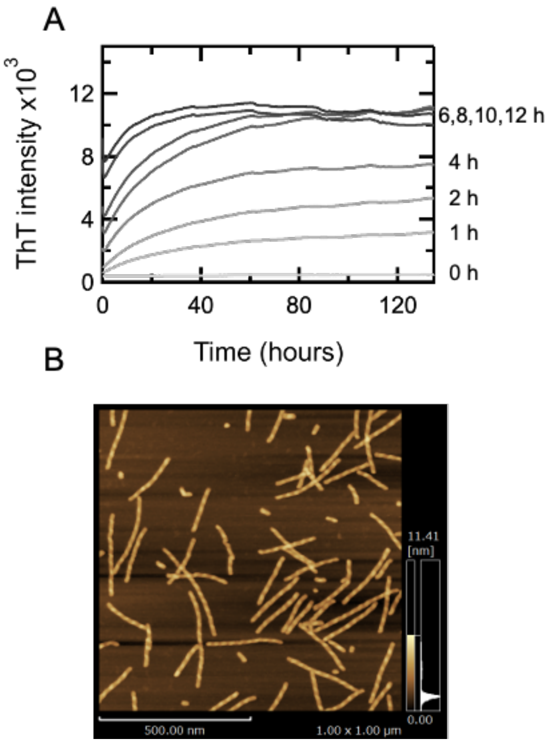


**Fig. S10. Shear stress confers fibrillogenic competence on GU under 3 M urea conditions.** (*A*) GU was diluted to 5 mg/mL in 3 M urea and subjected to shear by vortex mixing for the durations indicated, after which the time course of ThT fluorescence was recorded for approximately 130 h. Time zero corresponds to the start of the fluorescence measurement, that is, to the end of the shear treatment; the traces are therefore offset from one another by the duration of the treatment applied, and samples treated for longer periods already show elevated fluorescence at time zero. In the absence of shear (0 h), no increase in fluorescence was detected over the entire period. With shear, fluorescence increased, and the intensity at 130 h increased monotonically with the duration of treatment, approaching saturation between 6 and 12 h. (*B*) AFM image of GU after shear treatment for 6 h at 1,800 rpm (see below). Fibrils of uniform width are observed, together with dispersed nanoparticles. The fibrils exhibit periodic height modulation along the fibril axis, as observed for fibrils formed from UU (Fig. 2 A and C), and are shorter than the latter, consistent with shear both promoting fibril formation and fragmenting the fibrils formed. Scan area, 1.0 × 1.0 µm; height scale, 0–11.4 nm. For comparison, no structures are observed for GU in the absence of shear (Fig. 2B). ThT, thioflavin T; GU, guanidine–urea; AFM, atomic force microscopy.

**Shear treatment.** GU stock solution was diluted to 5 mg/mL with 3 M urea solution (10 mM Tris-HCl, 5 mM DTT, pH 7.0), and 300 µL aliquots were dispensed into 1.5 mL microcentrifuge tubes. Shear was applied with a vortex mixer (S0200, Labnet) operated at 3,400 rpm: the upper part of the tube was held by hand and its base was pressed against the centre of the mixer head so that the bottom of the tube described a circular path. Treatment was applied continuously at room temperature; no rise in sample temperature was detected. Treatment times were 0, 1, 2, 4, 6, 8, 10, and 12 h. Immediately after treatment, ThT was added and the fluorescence time course was recorded as described for the ThT kinetics above. The shear rate was not quantified. For AFM, a separate sample was treated for 6 h with a vortex mixer operated at 1,800 rpm, deposited on mica and imaged as described above.

**Tables**

**Table S1.** Secondary structure contents estimated using CDPro

| | α-helix (Regular) | α-helix (Distorted) | β-sheet (Regular) | β-sheet (Distorted) | Turn | Unstructured |
|---|---|---|---|---|---|---|
| UU | 0.1 | 2.4 | 8.5 | 5.9 | 10.1 | 72.5 |
| GU | 0.1 | 2.7 | 8.4 | 6.4 | 9.3 | 73.6 |

Values are presented in %. UU, urea–urea; GU, guanidine–urea.

**Table S2.** Secondary structure contents estimated using CDPro during fibril elongation

| | α-helix (Regular) | α-helix (Distorted) | β-sheet (Regular) | β-sheet (Distorted) | Turn | Unstructured |
|---|---|---|---|---|---|---|
| 0 h | 0.2 | 2.6 | 12.2 | 7.6 | 12.5 | 65.0 |
| 120 h | 0.2 | 2.8 | 18.4 | 9.8 | 17.4 | 51.4 |

Values are presented in %.

**Table S3.** Secondary structure contents estimated using BeStSel during fibril elongation

| | Helix1 (Regular) | Helix2 (Distorted) | Anti1 (left-twisted) | Anti2 (relaxed) | Anti3 (right-twisted) | Parallel | Turn | Others |
|---|---|---|---|---|---|---|---|---|
| 0 h | 0.0 | 0.0 | 0.0 | 12.0 | 15.8 | 0.0 | 13.7 | 58.1 |
| 120 h | 2.9 | 2.1 | 1.5 | 14.2 | 17.7 | 0.0 | 13.5 | 48.1 |

Values are expressed in %. Anti, antiparallel β-sheet.

**Table S4.** Results of peak deconvolution for the difference profile

| Peak | $2\theta$ (°) | | *d*-spacing (nm) | Direction |
|---|---|---|---|---|
| 1 | 16.1 | ±0.0235 | 0.55 | Equatorial |
| 2 | 18.8 | ±0.00116 | 0.47 | Meridional |
| 3 | 19.2 | ±0.00863 | 0.46 | Meridional |
| 4 | 23.0 | ±0.0834 | 0.39 | Meridional |

References


S1. S. C. Gill, P. H. von Hippel, Calculation of protein extinction coefficients from amino acid sequence data. *Anal. Biochem.* **182**, 319–326 (1989).

S2. N. Sreerama, R. W. Woody, Estimation of protein secondary structure from circular dichroism spectra: comparison of CONTIN, SELCON, and CDSSTR methods with an expanded reference set. *Anal. Biochem.* **287**, 252–260 (2000).

S3. A. Micsonai, *et al.*, BeStSel: a web server for accurate protein secondary structure prediction and fold recognition from the circular dichroism spectra. *Nucleic Acids Res.* **46**, W315–W322 (2018).

S4. B. Hammouda, Form factors for branched polymers with excluded volume. *J. Res. Natl. Inst. Stand. Technol.* **121**, 139–164 (2016).

S5. T. J. Prosa, B. J. Bauer, E. J. Amis, From stars to spheres: a SAXS analysis of dilute dendrimer solutions. *Macromolecules* **34**, 4897–4906 (2001).

**Movie S1 (separate file). HS-AFM movie of fibril growth performed under 1.7 M urea conditions**. The movie captures the elongation of fibrils from the precursor protein over a period of 549 s in a wide scanning area. Over time, both ends of the fibrils were observed to extend. This real-time observation supports the conclusion in the main text that fibril growth proceeds via the sequential polymerization (stepwise addition polymerization) of metastable dimers. The imaging was acquired with the continuous scanning mode at an imaging speed of 1.0 s/frame, and the video is played at 20× speed.
